# Caveolae respond to acute oxidative stress through membrane lipid peroxidation, cytosolic release of CAVIN1, and downstream regulation of NRF2

**DOI:** 10.1101/2021.06.09.447684

**Authors:** Yeping Wu, Ye-Wheen Lim, David A. Stroud, Nick Martel, Thomas E. Hall, Harriet P. Lo, Charles Ferguson, Michael T. Ryan, Kerrie-Ann McMahon, Robert G. Parton

**Affiliations:** The University of Queensland, Institute for Molecular Bioscience, 4072, Brisbane, Australia; The University of Melbourne, Department of Biochemistry and Pharmacology and The Bio21 Molecular Science and Biotechnology Institute, 3052, Parkville, Australia; Murdoch Children’s Research Institute, The Royal Children’s Hospital, 3052, Parkville, Australia; Monash University, Department of Biochemistry and Molecular Biology, Monash Biomedicine Discovery Institute, 3800, Melbourne, Australia; The University of Queensland, Centre for Microscopy and Microanalysis, 4072, Brisbane, Australia

**Keywords:** caveolae, CAVIN1, cell death, ferroptosis, lipid peroxidation, NRF2, oxidative stress, proteomics, wound healing

## Abstract

Caveolae have been linked to many biological functions, but their precise roles are unclear. Using quantitative whole cell proteomics of genome-edited cells, we show that the oxidative stress response is the major pathway dysregulated in cells lacking the key caveola structural protein, CAVIN1. *CAVIN1* deletion compromised sensitivity to oxidative stress in cultured cells and in animals. Wound-induced accumulation of reactive oxygen species and apoptosis were suppressed in Cavin1-null zebrafish, negatively affecting regeneration. Oxidative stress triggered lipid peroxidation and induced caveolar disassembly. The resulting release of CAVIN1 from caveolae allowed direct interaction between CAVIN1 and NRF2, a key regulator of the antioxidant response, facilitating NRF2 degradation. CAVIN1-null cells with impaired negative regulation of NRF2 showed resistance to lipid peroxidation-induced ferroptosis. Thus, caveolae, via lipid peroxidation and CAVIN1 release, maintain cellular susceptibility to oxidative stress-induced cell death demonstrating a crucial role for this enigmatic organelle in cellular homeostasis and wound response.

## Introduction

Caveolae are nanoscopic invaginations at the plasma membrane (PM) of most mammalian cells. Caveolar alterations have been extensively linked to human diseases while underlying mechanisms remain largely unknown^1^. Caveolae are generated by integral membrane proteins, caveolins, and cytoplasmic proteins, cavins. In the past, caveola studies focused on caveolins and their direct interaction with signalling proteins^2^ but these models have been questioned^3,4^. In recent years attention has turned to the newly discovered cavins^5–7^. These abundant structural proteins are increasingly linked to cancer^8–10^ and other disease conditions^11^ with different roles from caveolins^12,13^. Importantly, this demonstrates that caveolar function, and dysfunction in disease, cannot be understood without examining cavin roles.

Caveolae can flatten in response to increased membrane tension^14,15^. This results in a loss of caveolar structure, leading to a decreased association of caveolins and cavins^14,16,17^ and the redistribution of cavins into the cytosol^14,15,18^. Emerging evidence has shown that non-mechanical stimuli can also lead to cavin dissociation from caveolae. For example, insulin-activated signal can induce CAVIN1 release resulting in its targeting to the nucleus and regulation of ribosomal RNA synthesis in adipocytes^19^. Additionally, upon UV irradiation, CAVIN3 can be redistributed into the cytoplasm and nucleus to promote apoptosis^18^. Thus, we proposed that diverse cellular conditions and stimuli, such as oxidative stress that is ubiquitously associated with pathologies of tissues possessing caveolae^20^, can release cavins from the PM and lead to changes in intracellular targets [reviewed in^21^]. However, the upstream, downstream and the functional relevance of the cavin release-signalling model remains to be known.

To shed light on these questions, we investigated caveolar signaling by combining gene editing technology with unbiased whole cell quantitative proteomics and by the generation of zebrafish models to study the effect of Cavin1 loss *in vivo*. We showed that negative regulation of oxidative stress, mediated by nuclear factor-erythroid factor 2-related factor 2 (NRF2), was dramatically promoted in CAVIN1-null cells in culture and *in vivo*. The enhanced antioxidant capacity in CAVIN1-null cells and in zebrafish embryos led to oxidant-induced apoptosis resistance with *in vivo* consequences as demonstrated in zebrafish embryos. Wound-induced reactive oxygen species (ROS) and resulting apoptosis were suppressed in zebrafish lacking Cavin1, severely inhibiting the wound regeneration process. Although CAV1 has been previously linked to NRF2^22^, we show that CAV1 does not rescue the reduction of ROS in CAVIN1-null cells suggesting a novel pathway of oxidation regulation and/or sensing involving Cavin1. Mechanistic dissection in cultured cells showed that lipid peroxidation at the PM was required, and sufficient, to induce cytosolic redistribution of CAVIN1 under oxidative stress. Intracellular CAVIN1 released from caveolae was shown to interact with NRF2, promoting NRF2 degradation and triggering ferroptosis. Together, these data identify a novel pathway leading from lipid peroxidation at the plasma membrane to caveola disassembly and regulation of NRF2. In response to oxidative stress, CAVIN1 is released into the cytosol to negatively regulate NRF2 signalling, thereby maintaining cellular sensitivity to cell death, eliminating damaged cells and promoting regeneration after wounding.

## Results

### Cellular stress responses are impacted by CAVIN1 loss

We chose HeLa cells as a well-characterized model system for caveola studies^14,23,24^ . Wild-type (WT) HeLa cells express *CAV1*, *CAVIN1* and *CAVIN3* but not *CAVIN2* (Figure S1A). HeLa lines lacking CAVIN1 were generated by genome editing (Figure S1B-G) and further characterized by electron microscopy (EM) (Figure S1T), light microscopy (Figure S1M-S) and for protein and mRNA levels (Figure S1H-L).

Mass spectrometry (MS)-based quantitative proteomic analysis^25^ was performed to profile protein abundance and expression differences between WT and CAVIN1-null HeLa cells (Table S1). The significantly changed proteins in CAVIN1-null cells (Figure 1A; Table S1) were subjected to gene ontology and causal network analyses using Ingenuity Pathway Analysis (IPA) (Table S2, described in detail in the Supplementary Information [SI]). Cellular responses to stress were significantly affected by CAVIN1 loss (Table S2). Particularly, NRF2-mediated oxidative stress response was identified as the most upregulated toxicity pathway in CAVIN1-null cells (Figure 1B). Upstream regulator analysis (URA) further revealed NRF2, also known as NFE2L2, as a significant upstream regulator in CAVIN1-null cells (Figure 1C; entire URA list in Table S2).

**Figure 1.**
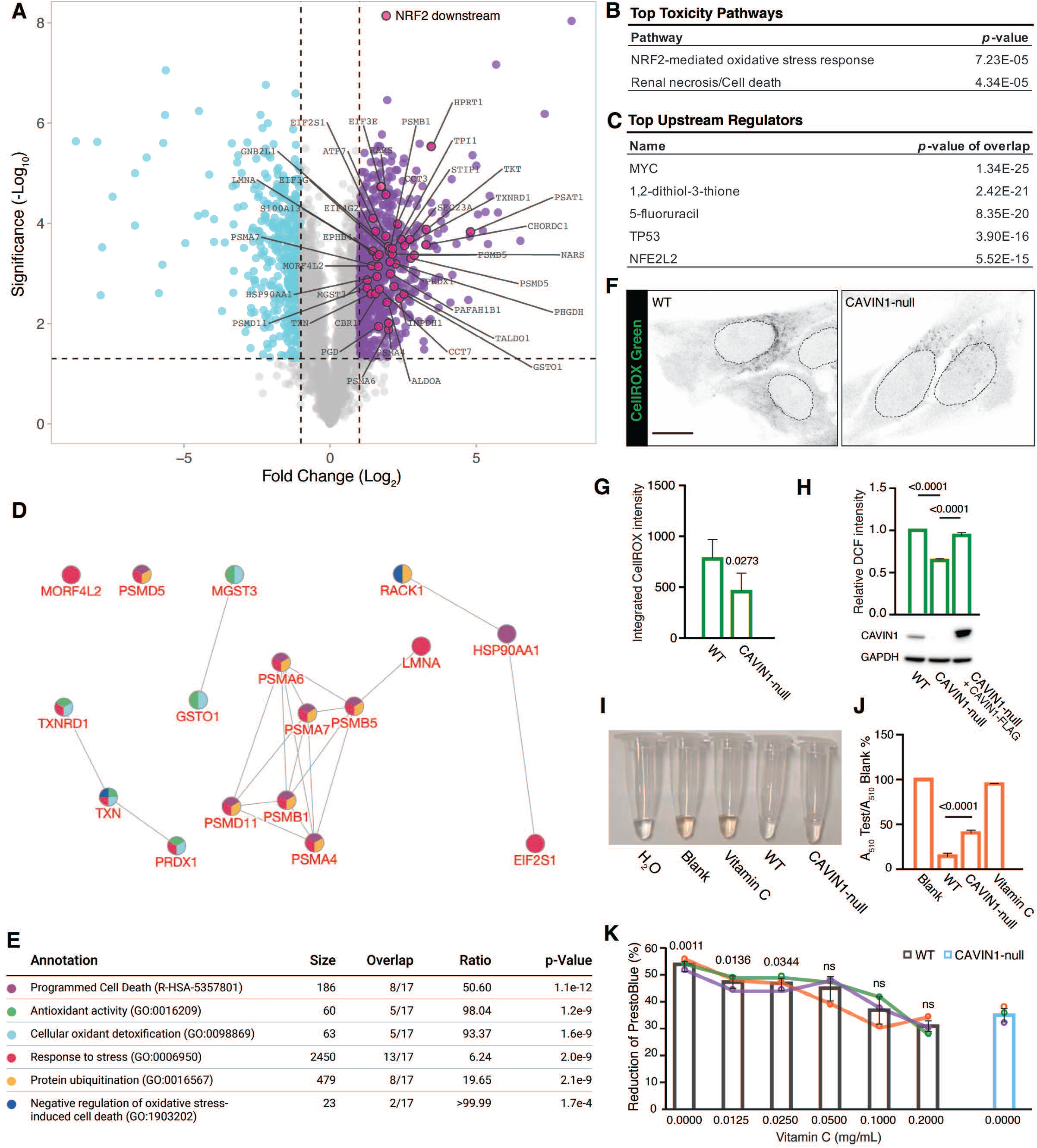
Comparative proteomics and pathway analysis reveal significant change in oxidative stress pathway in CAVIN1-null cells. (A) Volcano plot showing altered proteins CAVIN1-null cells relative to WT control. Horizontal dash line represents the 0.05 threshold on the p-values, while the vertical dash lines show threshold of ± 2 fold changes. Downregulated genes are indicated as cyan dots. Upregulated genes are colored in purple. Genes without significant change are labelled in gray. NRF2 target genes are highlighted in magenta. (B) Top toxicity pathways upregulated in CAVIN1-null cells. (C) Top upstream regulators responsible for upregulated pathways in CAVIN1-null cells, see entire list in Table S2. (D-E) Network of NRF2 downstream proteins (17/41). Solid lines indicate protein-protein interactions. Colored nodes present different biological processes as annotated in the enrichment table (E). (F) Inverted confocal images representing CellROX Green (5 μM, 30 min) stained cells. Circles indicate the outlines of nuclei. Scale bar, 10 μm. (G) Integrated intensity of CellROX Green fluorescence (two-tailed Student’s *t*-test, n=50 cells per experiment). (H) DCF intensity normalized to total protein amount (μg) in each group (one-way ANOVA with Tukey’s test). Western blot detected CAVIN1 levels in each sample with GAPDH as loading control. (I) Representative images of coloured-reaction samples. H_2_O was added as a negative control. (J) Absorbance was measured at 510 nm and compared using one-way ANOVA with Tukey’s test. (K) PrestoBlue assay was performed on cells after 48 h incubation with Vitamin C at different concentrations. Reduction of PrestoBlue (%) was statistically analyzed using one-way ANOVA with Dunnett’s test. Colored curve represents different experiment.

NRF2 is the key transcriptional regulator of antioxidants^26^. Under steady state conditions, NRF2 is rapidly degraded in the cytosol via the ubiquitin-proteasomal pathway^27^. Upon oxidative stress, NRF2 escapes degradation and translocates into the nucleus to activate downstream targets^26,28^. Although NRF2 was not detected in HeLa cells by MS, possibly due to its rapid turnover and low abundance at resting state conditions^27^, upregulated NRF2 in CAVIN1-null cells was verified by western blot analysis in whole cell lysates (Figure 3A-B). Forty-one significantly upregulated proteins were identified as NRF2 downstream targets in CAVIN1-null cells (highlighted in Figure 1A; Table 1). In general, these proteins are enriched in biological processes that negatively regulate oxidative stress (Figure 1D-E). For example, several proteins are antioxidant enzymes and negative regulators of oxidative stress (Figure 1D-E). In addition, proteins that constitute ubiquitin-proteasome system including proteasome subunit alpha type and beta type proteins (Figure 1D-E), have been shown to be essential for the proteolytic removal of oxidatively damaged proteins, enabling cells to cope with oxidative stress^29^. Together this suggests CAVIN1 depletion mediates an adaptive response to oxidative stress.

**Table 1.**
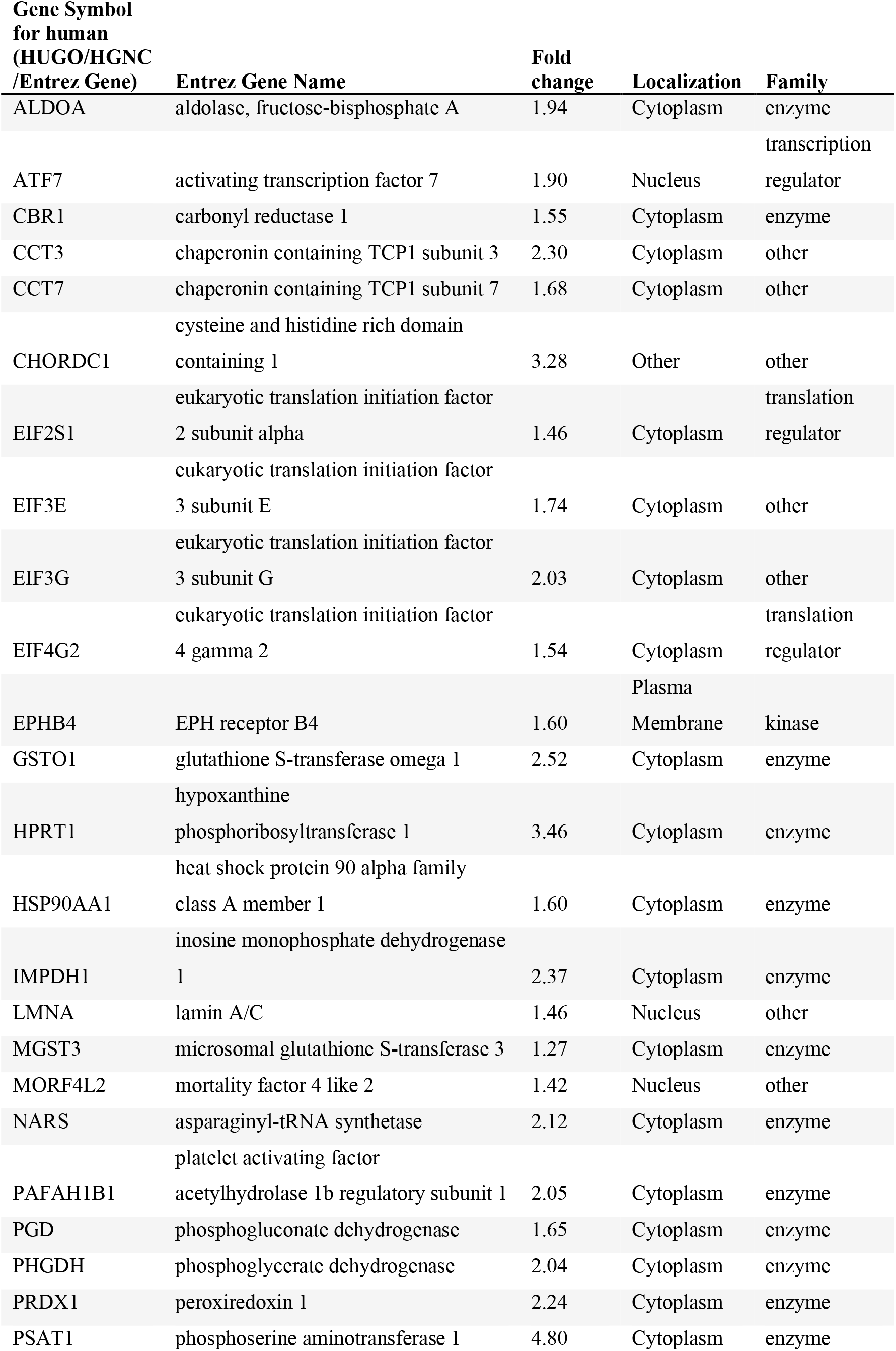
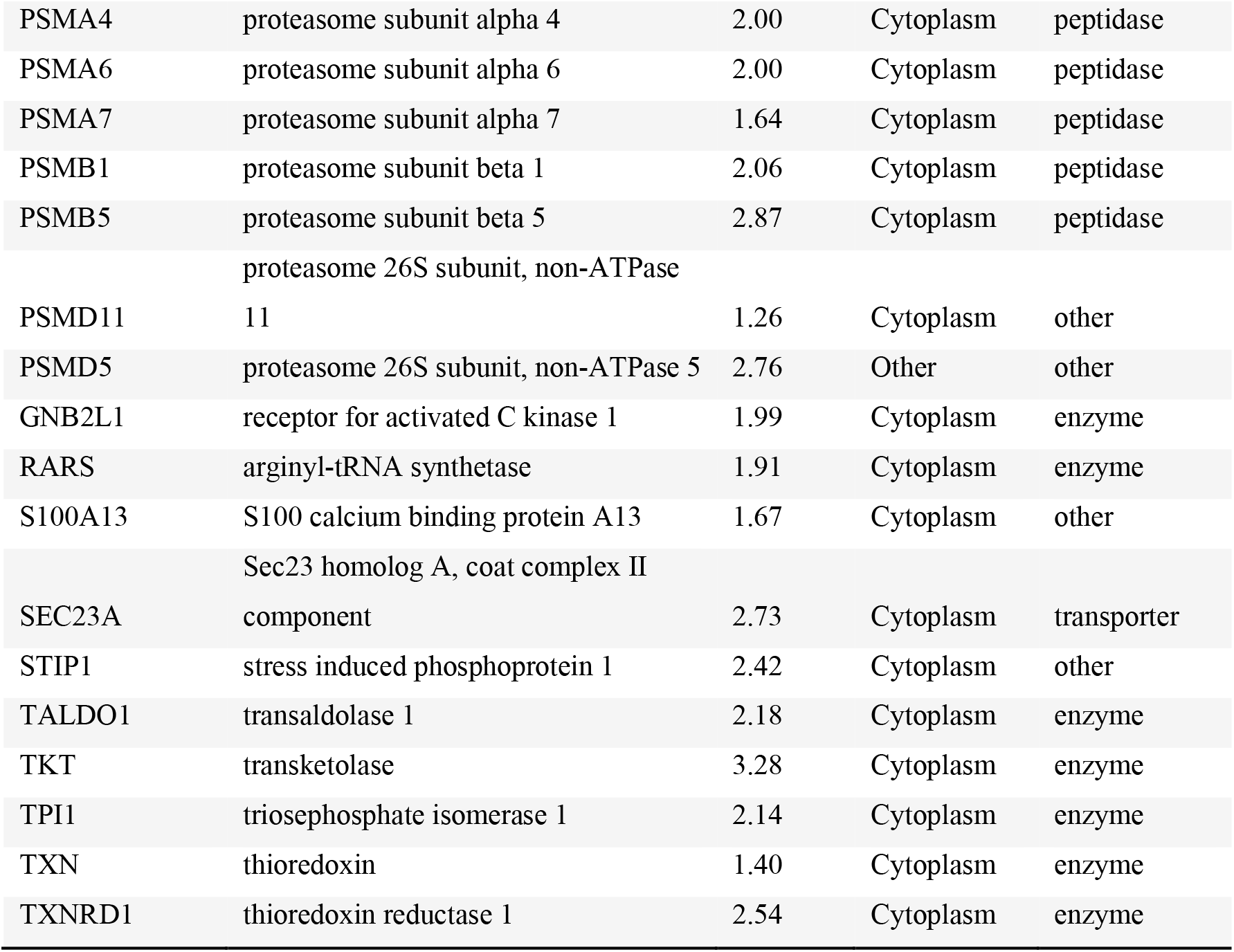
NRF2 downstream targets in CAVIN1-null HeLa cells identified by IPA.

### CAVIN1 ablation increases cellular antioxidant capacity

To elucidate the mechanism(s) underlying the upregulated antioxidant response observed in CAVIN1-null cells, we first assessed the antioxidant capacity of cells and zebrafish lacking CAVIN1.

Overall intracellular ROS content was measured using two probes; CellROXGreen and H_2_DCFDA. Both reagents verified a significant downregulation of basal ROS in CAVIN1-null cells (Figure 1F-H) that could be restored by CAVIN1-FLAG re-expression (Figure 1H). We also showed in an independent epidermoid carcinoma cell line, A431, that CAVIN1 knockdown led to reduced ROS levels (Figure S2A). Although CAV1 was downregulated in CAVIN1-null cells (Figure S1K-L), HA-CAV1 over-expression showed no effect on basal ROS levels (Figure S2B). This suggests that ROS reduction in CAVIN1-null cells under steady state conditions is a CAV1-independent effect. Next, a peroxide scavenging assay^30^ (Figure S2C) was performed with antioxidant Vitamin C as a positive control. CAVIN1-null cells showed significantly enhanced antioxidant capacity (Figure 1I-J), indicating upregulated ROS-detoxifying proteins in these cells.

Endogenous basal ROS as redox signaling messengers are needed for normal physiological function including cell proliferation^31^. With reduced ROS content, CAVIN1-null cells exhibited significantly downregulated cell growth ability (Figure S2D). We then treated WT cells with Vitamin C serial dilutions (Figure 1K) where the highest concentration (0.2 mg/mL) efficiently neutralized ROS (Figure S2E). Cell viability assay showed that Vitamin C suppressed WT cell proliferation in a dose-dependent manner (Figure 1K) where concentrations at or above 0.05 mg/mL abolished the difference from CAVIN1-null cells (Figure 1K). This suggests a correlation between suppressed cell growth and reduced basal ROS in CAVIN1-null cells.

### CAVIN1-null cells and zebrafish are oxidative stress-resistant

We next examined the cellular response to oxidative stress. Oxidant H_2_O_2_ was applied at a non-cytotoxic concentration (0.2 mM) (Figure S3H) for 30 min and ROS levels were monitored for the following 12 h (Figure 2A). After 1-h recovery, ROS levels showed no significant difference from stationary levels in CAVIN1-null cells. Excessive ROS was only effectively removed after a 6-h chase in WT cells (Figure 2A). Similar results were obtained with CellROX Green; ROS were restored to basal levels after 4 h in CAVIN1-null cells, while WT cells at this timepoint showed no significant reduction in ROS (Figure 2B-C). These data suggest upregulated ROS clearance ability in CAVIN1-null cells.

**Figure 2.**
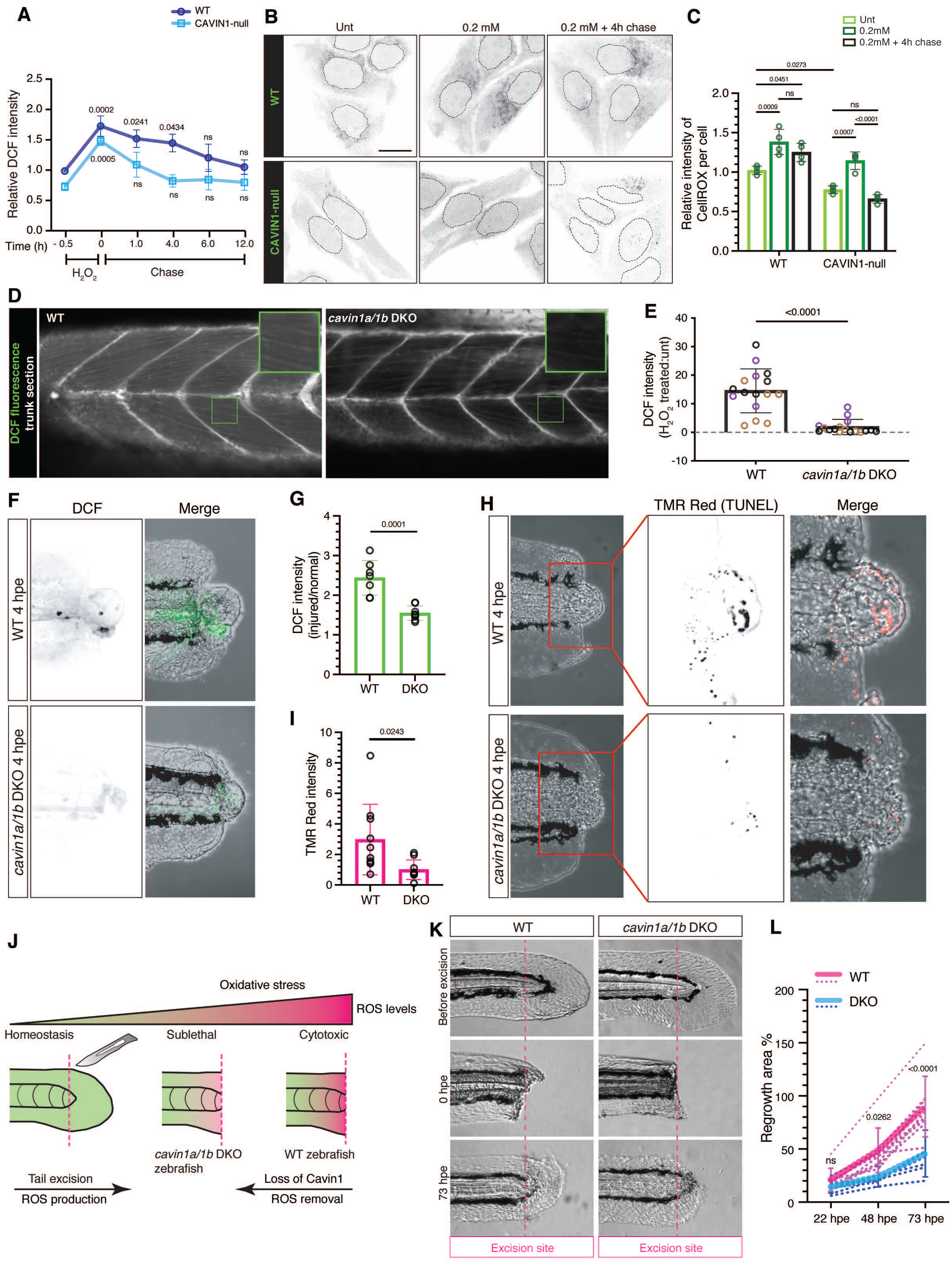
Loss of CAVIN1 leads to oxidative stress resistance. (A) Relative DCF fluorescence values in response to H_2_O_2_ treatment. Significant difference against values at -0.5 h was determined using two-way ANOVA with Sidak’s test. (B) Confocal images representing CellROX Green fluorescence in cells. Scale bar, 10 μm. (C) CellROX Green intensities were compared using two-way ANOVA with Tukey’s test. Colored dots represent different experiments. (D) Live confocal images displaying DCF fluorescence in skeletal muscle cells of 2 dpf WT and *cavin1a/1b* DKO zebrafish subjected to 2 mM H_2_O_2_ for 1 h. (E) Quantification and statistical analysis of DCF signal in H_2_O_2_ treated WT and *cavin1a/1b* DKO zebrafish normalized with DCF signal of clutch-matched untreated controls (two-tailed Student’s *t*-test, n=16 WT zebrafish and n=17 *cavin1a/1b* DKO zebrafish, 3 clutches per line). Colored dots represent different clutches. (F) Confocal images showing DCF signal at the wound site in 2 dpf WT and *cavin1a/1b* DKO zebrafish at 4 hpe. (G) DCF signal in the injured tissue was normalized to DCF signal in the normal tissue within the same zebrafish (two-tailed Student’s *t*-test, n=7 WT zebrafish and n=8 *cavin1a/1b* DKO zebrafish, 3 clutches per line). (H) TUNEL-detected apoptotic cells in 2 dpf WT and *cavin1a/1b* DKO zebrafish at 4 hpe. (I) Mean TMR-Red signal (to mean intensity of WT) at the wound site, indicating apoptotic levels in WT and *cavin1a/1b* DKO zebrafish (two-tailed Student’s *t*-test, n=10 WT zebrafish and n=9 *cavin1a/1b* DKO zebrafish, 3 clutches per line). (J) A diagram illustrating responses to tissue injury in WT and *cavin1a/1b* DKO. (K) Tail size before excision and at 0 and 73 hpe in WT and *cavin1a/1b* DKO. (L) Tail size at 22, 48 and 73 hpe was compared to the tail size before excision in the same zebrafish. Regrowth area % in WT and *cavin1a/1b* DKO zebrafish at each time point were compared using two-way ANOVA with Sidak’s test (n=8 WT zebrafish and n=8 *cavin1a/1b* DKO zebrafish, 3 clutches per line).

We then tested whether increased antioxidant capacity is an evolutionarily-conserved feature of tissues lacking CAVIN1. For this, we used the zebrafish as a vertebrate model system. The zebrafish expresses two Cavin1 paralogs, Cavin1a and Cavin1b^5,32^. To generate a complete Cavin1 double knockout (DKO) zebrafish line, hereby referred to as the *cavin1a/1b* DKO line, we generated a *cavin1a* CRISPR/Cas9 KO line (*cavin1a^-/-uq10rp^*) and crossed this line to our previously characterized *cavin1b* KO line (*cavin1b^-/-uq7rp^*)^33^; offspring were bred to homozygosity and characterized (Figure S3A-G, described in details in the SI). *Cavin1a/1b* DKO zebrafish lacked caveolae in all tissues examined (Figure S3G). We then investigated the ROS scavenging ability of *cavin1a/1b* DKO zebrafish. Live embryos (2 days post-fertilization [dpf]) were subjected to H_2_O_2_ and then stained with H_2_DCFDA for ROS detection. The myotome occupies a large proportion of the zebrafish trunk, therefore we focused on the DCF signal in skeletal muscle cells (Figure 2D). While H_2_O_2_-treated WT zebrafish showed significantly increased ROS, H_2_O_2_-treated *cavin1a/1b* DKO zebrafish showed similar ROS levels to untreated *cavin1a/1b* DKO zebrafish (Figure 2D-E). These *in vivo* results suggest that increased antioxidant capacity is a conserved feature of animals lacking Cavin1.

### CAVIN1-mediated ROS is linked to in vivo wound healing

Tissue damage can result in immediate ROS elevation to excessive levels, triggering apoptosis to limit detrimental effects of oxidative damage^34–36^. To assess the pathophysiological significance of Cavin1-mediated ROS regulation, we performed tail excision^37^ to induce acute injury and measured ROS and apoptotic levels in *cavin1a/1b* DKO zebrafish. ROS was induced at the wound site at 4 h post excision (hpe) (Figure 2F). *Cavin1a/1b* DKO zebrafish showed significantly decreased ROS accumulation at the wound site (1.5-fold) compared to WT zebrafish (2.4-fold) (Figure 2F-G). TUNEL revealed that cells at the wound site in WT zebrafish underwent apoptosis at 4 hpe. This was significantly suppressed in *cavin1a/1b* DKO zebrafish (Figure 2H-I). This shows that with enhanced buffering ability, wound-induced ROS could be maintained at sublethal levels in *cavin1a/1b* DKO zebrafish which allows the survival of damaged cells (Figure 2J). Interestingly, previous studies revealed an unexpected role of ROS and apoptotic signal in the stimulation of regenerative proliferation^38–40^. By examining tail regeneration, we found that regrowth tail area (%) in *cavin1a/1b* DKO zebrafish was decreased from 22 to 73 hpe, significantly different from WT zebrafish at 48 and 73 hpe (Figure 2K-L). Together, this suggests that Cavin1 ablation impairs wound-healing process in zebrafish by suppressing ROS and apoptosis.

Complementing these results, cell viability assays revealed significantly increased survival rates of CAVIN1-null cells treated with H_2_O_2_ at toxic levels (Figure S3H). Reduced levels of DNA damage marker γH2AX and apoptotic marker cleaved caspase 3, were also observed in H_2_O_2_-treated CAVIN1-null cells (Figure S3I). Collectively, these results show that CAVIN1 deletion enhances antioxidant capacity, leading to resistance to oxidative stress-induced cell death.

### CAVIN1 regulates NRF2 nuclear import, function and degradation

Having confirmed increased antioxidant capacity in cells and animals lacking CAVIN1, we next examined if this effect could be a result of changes in the NRF2 pathway (Figure 1B-C). Increased NRF2 in CAVIN1-null cells was detected under steady state conditions (Figure 3A-B). After H_2_O_2_ exposure, significant NRF2 upregulation was observed in CAVIN1-null cells at 40 min, while NRF2 was upregulated, but not significantly, at 80 min in WT cells (Figure 3A-B). Immunofluorescence confirmed significantly increased NRF2 nuclear import in CAVIN1-null cells under both steady state and oxidative stress conditions (Figure 3C-D). CAVIN1-GFP re-expression retained NRF2 in the cytoplasm in both untreated and H_2_O_2_-treated CAVIN1-null cells (Figure 3E), which was not observed in untransfected (Figure 3E, white arrows) or GFP-transfected cells (Figure 3F). In the CAVIN1/caveola-deficient MCF-7 cell line^18^, NRF2 exhibited primarily in the nuclei (Figure S4A). In these cells, CAVIN1-GFP re-expression also induced cytosolic redistribution of NRF2 (Figure S4A). These data suggest that CAVIN1 mediates cytosolic sequestration of NRF2.

**Figure 3.**
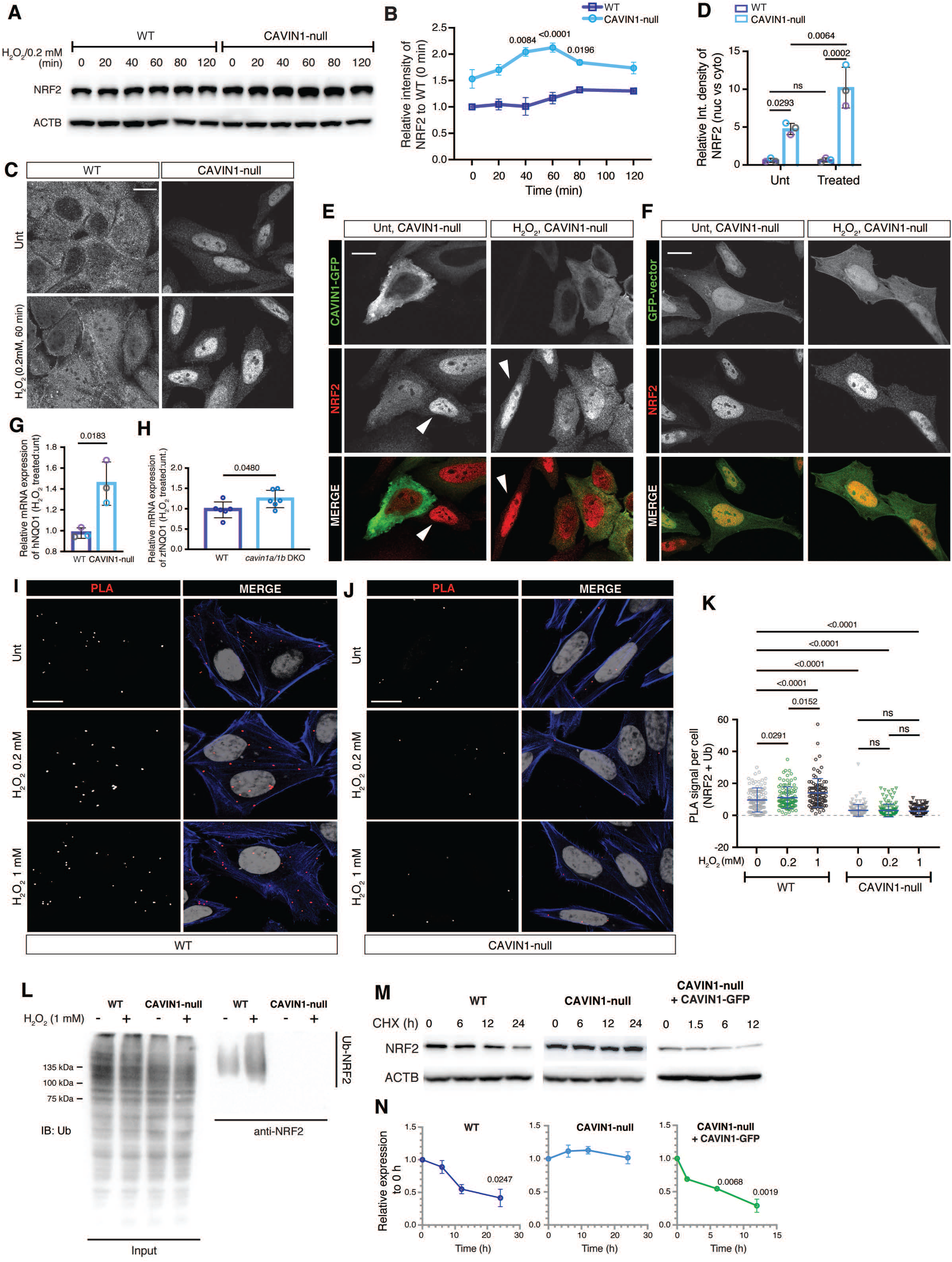
CAVIN1 is essential for the ubiquitination of NRF2 under oxidative stress. (A) Western blotting analysis of NRF2 protein expression following 0.2 mM H_2_O_2_ treatment over a 120-min time course. (B) Relative NRF2 intensity values in (A) were then calculated and statistically compared against NRF2 intensity at 0 min for both WT and CAVIN1-null cells (two-way ANOVA with Sidak’s test). (C) NRF2 immunofluorescence in untreated and H_2_O_2_-treated WT HeLa cells. Scale bar, 10 μm. (D) The ratio of nuclear NRF2 intensity to cytoplasmic NRF2 intensity (nuc vs. cyto) was calculated (n=50 cells per experiment) and compared using two-way ANOVA with Tukey’s test. (E-F) NRF2 immunostaining in CAVIN1-null cells transfected with CAVIN1-GFP (E) or GFP-vector (F) and treated with H_2_O_2_ (0.2 mM, 60 min). White arrows: untransfected CAVIN1-null cells. Scale bar, 10 μm. (G-H). Real-time PCR was performed to assess the fold change (H_2_O_2_ treated:untreated) of NQO1 transcripts in CAVIN1-null cells (G) and in *cavin1a1b* DKO zebrafish (H) following 80 min H_2_O_2_ treatment. TBP was used as a housekeeping gene. Statistical difference was assessed by two-tailed Student’s *t*-test. (I-J) Representative images showing PLA signal of NRF2-ubiquitin association in WT (I) and CAVIN1-null (J) cells. MG132 (10 μM) was applied to cells for 3 h prior to H_2_O_2_ treatment. (K) Quantification and statistical analysis of PLA signal (n=50 cells per experiment, two-way ANOVA with Tukey’s test). (L) Cells were treated with MG132 at 10 μM for 3 h and were then subjected to 1 mM H_2_O_2_ treatment for 60 min. Cell lysates were harvested for NRF2 immunoprecipitation and western blot assays. Ubiquitin was examined in NRF2 pull-down samples. (M) Western blotting analysis of NRF2 levels following CHX (50 μg/mL) treatment. (N) Relative NRF2 levels in CHX chase assays was calculated and statistically analyzed using one-way ANOVA with Dunnett’s test.

Next, we assessed NRF2 function as a transcriptional factor. The antioxidant enzyme and prominent NRF2 target, NQO1^41^, was upregulated in our proteomic analysis (Table S1); real-time PCR demonstrated significantly increased NQO1 mRNA levels in H_2_O_2_-treated CAVIN1-null cells and *cavin1a/1b* DKO zebrafish (Figure 3G-H). These results confirmed increased transcriptional activity of NRF2 in CAVIN1-null cells and zebrafish.

Cytosolic NRF2 is rapidly degraded via the ubiquitin-proteasome pathway^42^. We next asked whether CAVIN1-mediated cytosolic sequestration of NRF2 affects its protein stability. First, we assessed NRF2 ubiquitination levels. Proximity ligation assay (PLA) (Figure 3I-K) and co-IP (Figure 3L) verified decreased NRF2-ubiquitin association in both untreated and H_2_O_2_-treated CAVIN1-null cells, suggesting the essential role of CAVIN1 in mediating NRF2 ubiquitination. To evaluate protein stability, NRF2 was examined over a 24-h period of cycloheximide (CHX) treatment (Figure 3M-N). After a 24-h chase, significant reduction of NRF2 was observed in WT cells. In contrast, no obvious NRF2 downregulation was observed in CAVIN1-null cells within 24 h. CAVIN1-GFP re-expression strongly promoted NRF2 degradation in CAVIN1-null cells, verifying that CAVIN1 is directly linked to NRF2 stability.

### CAVIN1 is released from caveolae and interacts with NRF2 in the cytosol under oxidative stress

CAVIN1 as a caveola-associated protein is mainly localized on the cell surface^5^. NRF2 is cytoplasmic under steady state conditions (although a surface pool was also detectable) (Figure 3C). The difference in subcellular localization for these two proteins raises the question of how CAVIN1 regulates NRF2 activity in WT cells. As intracellular redistribution of CAVIN1 has been observed under different stress conditions previously^5,14,15,18,19^, we examined whether oxidative stress would also induce CAVIN1 release. In H_2_O_2_-treated WT HeLa cells, CAVIN1 was redistributed from the cell surface (Figure 4A) to the cytosol (Figure 4B-C). Similar observations were obtained in A431 cells (Figure S4B). This was correlated with a decreased density of surface caveolae detected by EM (Figure S4C-E). Live-cell imaging further revealed reduced CAV1-CAVIN1 colocalization and cytosolic redistribution of CAVIN1 in the H_2_O_2_-treated WT cell (Figure 4D-I), suggesting dissociation of CAVIN1 from caveolae. Next, we observed significantly upregulated CAVIN1-NRF2 association in H_2_O_2_-treated WT cells in a dose-dependent manner using PLA (Figure 4J-K) and co-IP (Figure 4L-P). Notably, PLA showed that this increased association is cytosolic (Figure 4J). Collectively, these data demonstrate that CAVIN1 release is a general effect upon oxidative stress, allowing CAVIN1 to functionally associate with NRF2 in the cytosol.

**Figure 4.**
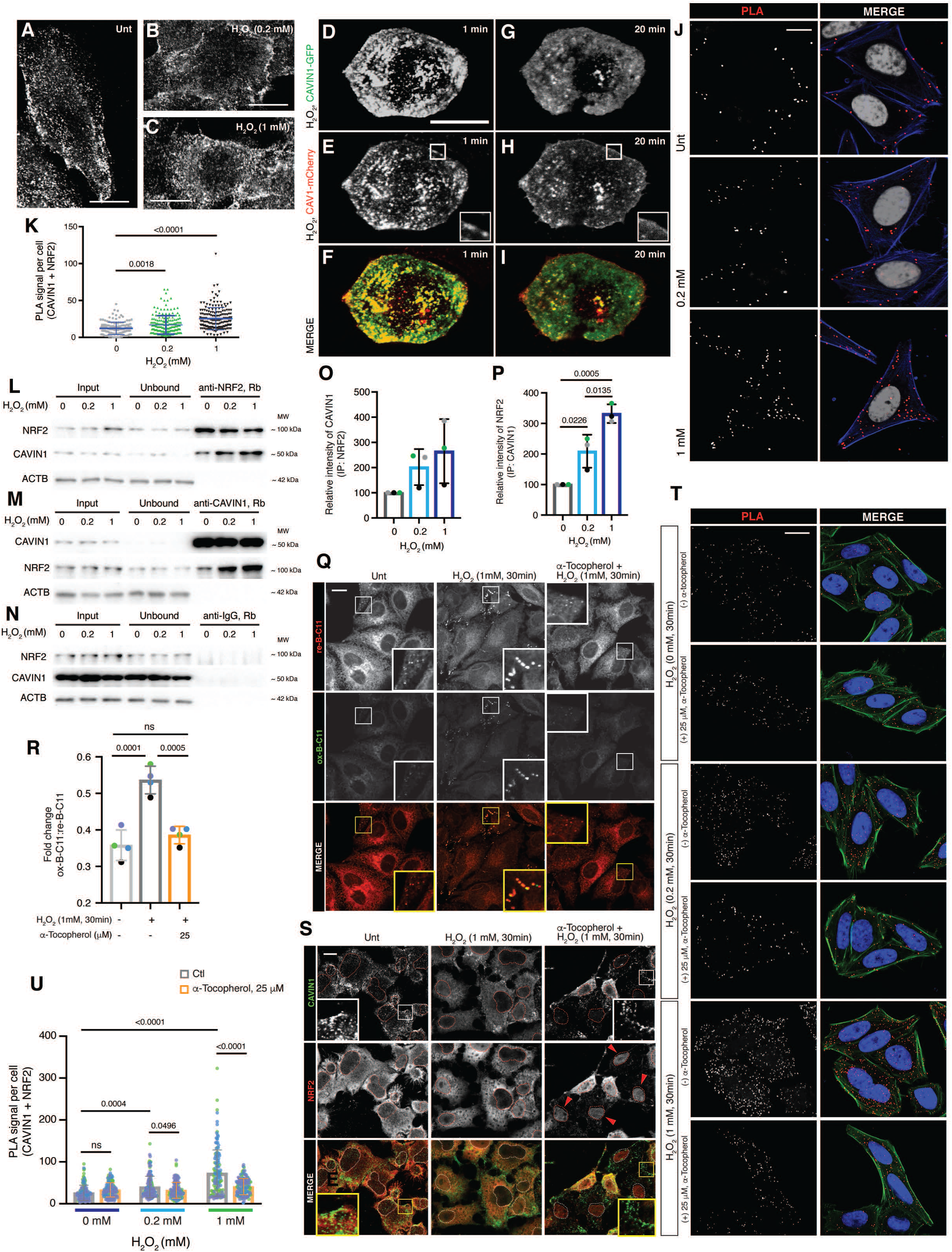
LPO causes CAVIN1 release from caveolae and the interaction between CAVIN1 and NRF2 in the cytoplasm. (A-C) Representative confocal images showing CAVIN1 distribution in untreated (A) or H_2_O_2_ treated (B-C) WT HeLa cells. Scale bar, 10 μm. (D-I) Live cell imaging of WT HeLa cells co-transfected with CAVIN1-GFP and CAV1-mCherry with 1 mM H_2_O_2_ treatment. Insets shows enlargement of the selected areas. A diffuse pattern of CAV1 at the PM (H) was used as an indicator of partial caveola disassembly^6^. Scale bar, 10 μm. (J) Images of PLA signal showing the CAVIN1-NRF2 association before and after 60-min H_2_O_2_ treatment. PLA signal alone and DAPI staining in merged images were inverted to grey scale. Phalloidin staining (blue) labels the boundary of the cells. Scale bar, 10 μm. (K) PLA signal per cell in (J) was statistically analyzed using one-way ANOVA with Tukey’s test. (L-N) Immunoprecipitation assays assessed the interaction between NRF2 and CAVIN1 upon 60 min H_2_O_2_ treatment in both anti-NRF2 pull-down samples (L) and anti-CAVIN1 pull-down samples (M). Anti-IgG pull-down samples were assessed as negative controls (N). (O-P) Quantification and statistical analysis of CAVIN1 levels (O) or NRF2 levels (P) in pulled down samples using one-way ANOVA with Tukey’s test. Colored dots represent different experiments. (Q) Confocal images showing LPO in WT cells using B-C11 probes. Scale bar, 10 μm. (R) Fold changes of ox-B-C11:re-B-C11 were statistically analyzed by one-way ANOVA with Tukey’s test. (S) Immunofluorescence assays showing the effect of α-Tocopherol (25 μM) on the distribution of CAVIN1 (green) and NRF2 (red). Nuclei are indicated as red circles in the inverted grey scale images and as white circles in the merged images. Scale bar, 10 μm. (T) PLA detected the CAVIN1-NRF2 association under H_2_O_2_ treatment in WT cells with or without pre-treatment by α-Tocopherol (25 μM). Scale bar, 10 μm. (U) PLA signal in (T) was quantified and compared using one-way ANOVA with Tukey’s test.

### Lipid peroxidation causes CAVIN1 release upon oxidative stress

It has been previously suggested that CAVIN1 is associated with the PM through electrostatic interactions with phosphoinositide (PI) lipids^43^ and phosphatidylserine^44^ and we have proposed that specific lipid species may be enriched in caveolae^45^. Lipid peroxidation (LPO) has been considered as a main cause of PM damage under oxidative stress^46^. Therefore, we hypothesized LPO as a potential mechanism for caveola disassembly and CAVIN1 redistribution. To test this, we used a live-cell LPO sensor, BODIPY-C11 (B-C11)^47^. Significantly increased (oxidized B-C11 [ox-B-C11]:reduced B-C11 [re-B-C11]) ratio in H_2_O_2_-treated cells indicated LPO occurrence (Figure 4Q-R). Cumene hydroperoxide, a well-characterized LPO inducer^48^, was included as a positive control (Figure S4I-J). Pre-treatment with lipid soluble antioxidant α-Tocopherol that is protective against LPO^49^ restored (ox-B-C11:re-B-C11) ratio to basal levels in H_2_O_2_-treated cells (Figures 4Q-R). Similar results were observed with A431 cells (Figure S4G-H).

Having optimized LPO detection methods, we next asked whether LPO is involved in CAVIN1 release upon oxidative stress. Immunofluorescence revealed that CAVIN1 redistribution and NRF2 cytosolic sequestration were inhibited by α-Tocopherol in H_2_O_2_-treated WT cells (Figure 4S), in which NRF2 efficiently translocated to the nucleus (Figure 4S, red arrows). Treatment with α-Tocopherol also abrogated the H_2_O_2_-induced increase in CAVIN1-NRF2 association and restored the interaction signal at the PM (Figure 4T-U).

We next tested whether LPO is sufficient to induce CAVIN1 release. Cumene hydroperoxide induced CAVIN1 release and this was blocked by α-Tocopherol (Figure S4K). Moreover, we used a direct photochemical method to bypass enzymatic redox reactions and specifically initiate LPO at the PM^50^. Live-cell imaging with confocal microscopy revealed that 405 nm high-power laser pulses applied on the PM led to increased ox-B-C11 signal compared to the untreated cell (Video S1-2; Figure S5A-B). We then used this treatment on monolayer regions containing multiple cells and stained endogenous CAVIN1. Cells in the laser-treated region exhibited less CAVIN1 PM puncta (Figure 5A-B) and increased ox-C-B11 signal (Figure 5C-D) compared to the cells in the untreated region (Figure 5A and 5I). α-Tocopherol inhibited LPO (Figure 5G-H) and the loss of CAVIN1 puncta at the PM (Figure 5E-F and 5I). Live-cell imaging showed decreased CAVIN1-GFP puncta at the PM over time in the laser-treated cell compared to the untreated cell (Figure S5C-F). These results implicate LPO at the PM in triggering caveolar disassembly and CAVIN1 release.

**Figure 5.**
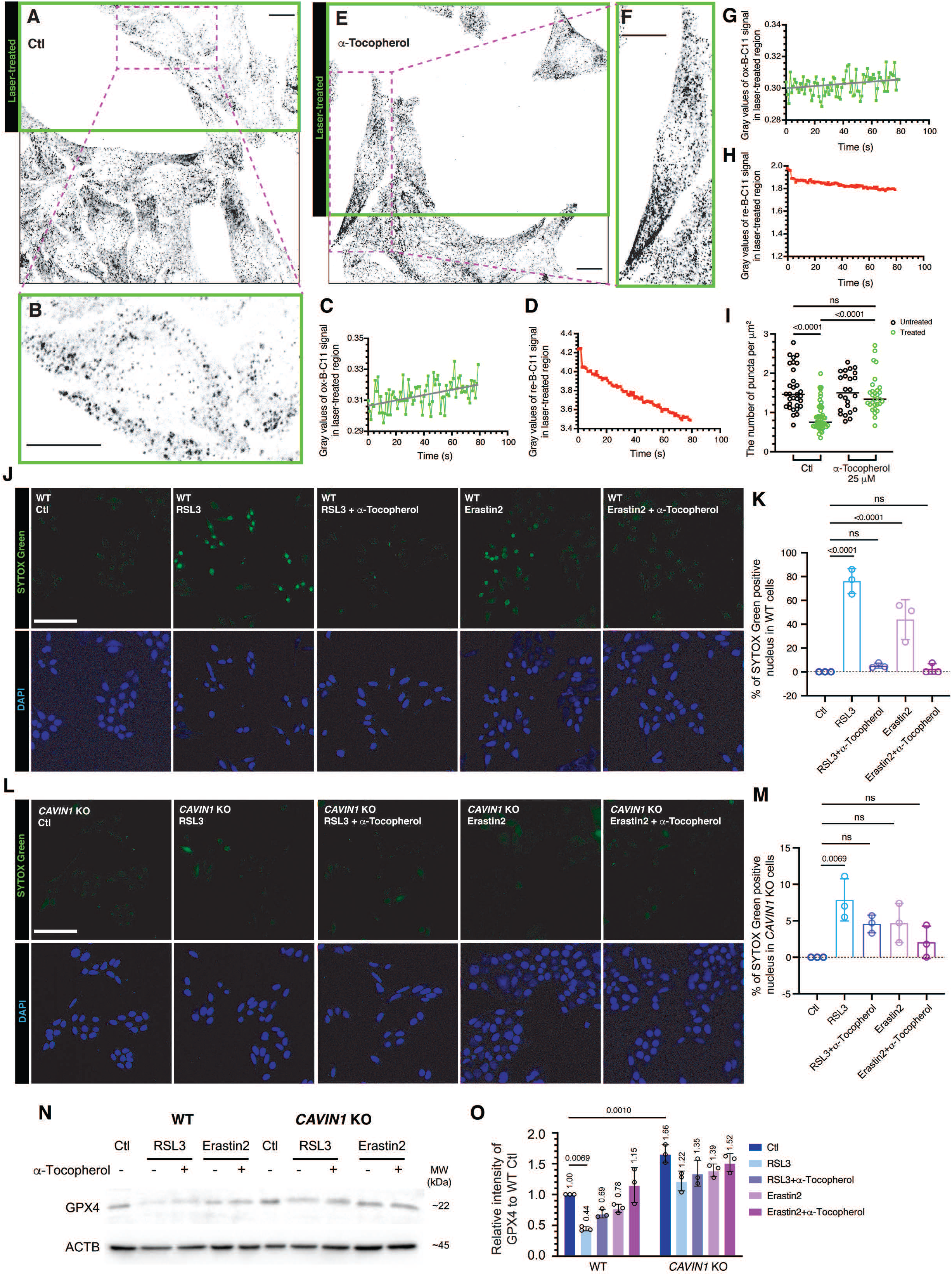
Peroxidation of membrane lipids induces CAVIN1 dissociation from caveolae and leads to ferroptosis. (A) Inverted confocal images showed the density of endogenous CAVIN1 puncta at the PM in WT HeLa cells. In the selected region (green frame), cells were subjected to 405 nm laser pulses at the PM. Scale bar, 10 μm. (B) Enlarged image showing the number of CAVIN1 puncta at the PM in the laser-treated cell. Scale bar, 10 μm. (C-D) Gray values of ox-B-C11 signal (C) and re-B-C11 (D) in the treated region were quantified to over 80 sec following laser treatment. (E) Immunofluorescence showing endogenous CAVIN1 puncta in cells pre-treated with α-Tocopherol. Green frame indicates the region subjected to 405 nm laser pulses. Scale bar, 10 μm. (F) Enlarged image showing CAVIN1 puncta at the PM in the laser-treated cell in the presence of α-Tocopherol. Scale bar, 10 μm. (G-H) Oxidized (G) and reduced (H) B-C11 signal were quantified for the treated region presented in (E). (I) The number of CAVIN1 puncta at the PM in untreated and laser-treated cells with or without pre-treatment by α-Tocopherol (25 μM) were compared using two-way ANOVA with Tukey’s test. (J-M) SYTOX Green staining revealed ferroptotic WT (J-K) or CAVIN1-null (L-M) cells following 24-h RSL3 (5 μM) or Erastin2 (5 μM) treatment with or without pre-treatment by α-Tocopherol (25 μM). The percentage (%) of SYTOX Green positive cells were calculated and statistically analyzed using one-way ANOVA with Tukey’s test. DAPI stain was used for the indication of total number of cells in each frame. Scale bar, 10 μm. (N) GPX4 levels were detected by western blot assays. (O) Relative GPX4 levels were calculated, and significant change was indicated by two-way ANOVA with Sidak’s test.

### CAVIN1-null cells are resistant to ferroptosis

Oxidative stress can induce several types of cell deaths including LPO-dependent ferroptosis^51,52^. NRF2 has been shown to be a negative regulator of LPO and ferroptosis^53^. In view of the upregulated NRF2 signaling in CAVIN1-null cells, we therefore assessed the sensitivity of these cells to ferroptosis induced by RSL3^54,55^ and Erastin2^52,56,57^. Cells undergoing ferroptosis display positive staining of cell impermeable SYTOX Green nucleic acid dye^56^. As indicated by SYTOX Green staining, ferroptosis were effectively induced in RSL3- or Erastin2-treated WT cells (Figure 5J-K). This was blocked by α-Tocopherol (Figure 5J-K). Strikingly, decreased percentage of dead cells was observed in CAVIN1-null cells (7.8%) compared to WT cells (76.2%) after RSL3 treatment (Figure 5J-M). Erastin2 sensitivity was also decreased in CAVIN1-null cells (Figure 5L-M). Western blot further revealed upregulated glutathione peroxidase 4 (GPX4), a key phospholipid hydroperoxidase^58^, in CAVIN1-null cells at steady state conditions (1.66-fold) (Figure 5N-O). As a GPX4 inhibitor, RSL3 induced significant downregulation of GPX4 (−56%) in WT but not CAVIN1-null cells (−26%) (Figure 5N-O). This RSL3-induced GPX4 reduction was rescued by α-Tocopherol. These results show resistance to ferroptosis in cells lacking CAVIN1.

Altogether, our study showed that LPO under oxidative stress causes CAVIN1 release from caveolae, providing a mechanism by which CAVIN1 can interact with NRF2 to sequester it in the cytosol and promote its degradation. This subsequently leads to cell death such as ferroptosis upon severe oxidant attack (schematic conclusion in Figure 6).

**Figure 6.**
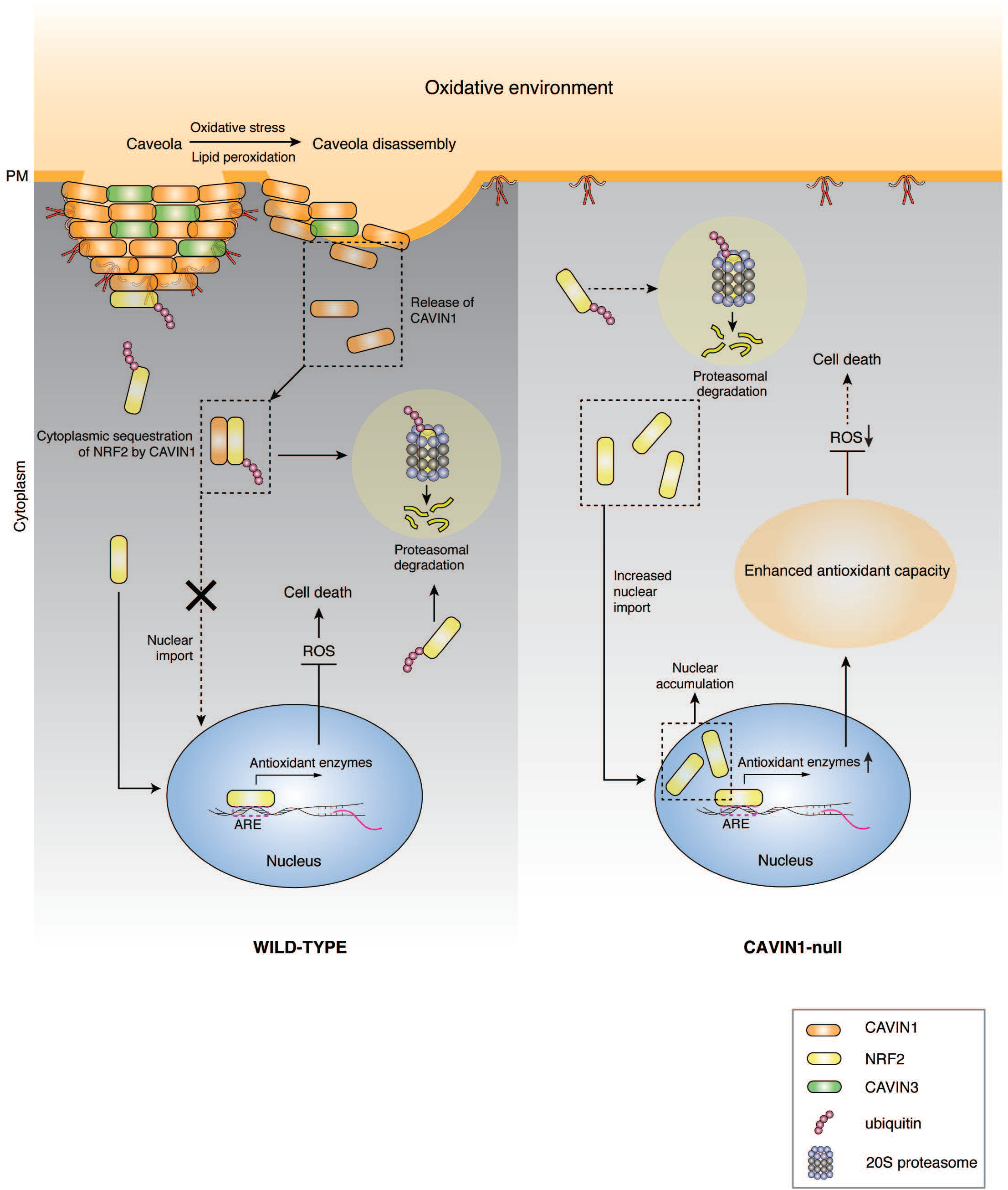
A graphical summary illustrating the molecular role of CAVIN1 in response to oxidative stress. Oxidative stress-induced LPO leads to CAVIN1 release from caveolae to the cytosol. This is essential for the sequestration and degradation of NRF2 and leads to cell deaths. In contrast, loss of CAVIN1 allows a more efficient NRF2 nuclear accumulation and up-regulates NRF2 downstream antioxidant enzymes upon oxidative stress. This confers enhanced antioxidant capacities to the cells and resistance to excess ROS-induced cell cytotoxicity.

## Discussion

Our results have implications for understanding both the effect of caveola loss in cancer cells and the physiological role of caveolae as part of the cellular response to stress. First, we show via unbiased proteomics that one of the most striking effects of caveola loss, which can occur in cancer cells with abnormal expression of CAVIN1^5,12,18^, is the upregulation of antioxidant activity due to impaired negative regulation of the NRF2 pathway. This promotes insensitivity to oxidative stress-induced cell death such as ferroptosis, increasing the survival of cells with malignant potential. Second, we propose a scenario in which caveolae can regulate the cellular response to extreme oxidative stress (Figure 6), as demonstrated with an acute tissue injury and regeneration assay.

Under steady state conditions, certain levels of ROS are required for normal cellular functions such as cell proliferation^31^. In our study, CAVIN1-null cells with reduced basal ROS levels showed compromised cell growth, indicating that caveolae play an important role in maintaining ROS-induced signaling in healthy cells. Under oxidative stress, which can be induced during pathophysiological conditions^38,59^, ROS can be elevated to lethal levels. We propose that caveolae, due to their unique lipid composition^45^ are particularly sensitive to peroxidation of key unsaturated fatty acid chains on membrane lipids. The resulting bilayer changes caused by LPO^46^ leads to caveola disassembly and the dissociation of CAVIN1 from the PM to interact with cytoplasmic NRF2, facilitating NRF2 ubiquitination and degradation. This triggers ferroptosis, a recently recognized LPO-driven form of cell death^52^. During cancer progression, increased expression of negative regulators of oxidative stress such as NRF2 can suppress ferroptosis and promote therapy resistance^53,60,61^. Our data show that cancer cells lacking caveolae exhibit upregulated NRF2 and are ferroptosis resistant. This suggests that prospectively, CAVIN1/caveola expression level may serve as a novel indicator of chemosensitivity, directing selection of patients responsive to ferroptosis-related cancer therapies^60,62^.

Using a well-characterized wound assay in the zebrafish, we showed that *cavin1* loss decreased injury-induced ROS accumulation, and this was associated with reduced apoptosis (Figure 2J). Activation of an apoptotic signal is required for both damaged cell removal and propagation of wound repair signals required for healing^40^. Consistent with this healing was significantly impaired in the *cavin1/1b* DKO embryos. A sufficient level of wound-induced extracellular H_2_O_2_ has also been shown to be required in the recruitment of immune cells and Src family kinase members during regeneration^63–65^, suggesting that the epimorphic regeneration delay observed in amputated *cavin1/1b* DKO embryos may involve changes in a more complex system involving both redox and apoptotic cell signalling.

CAVIN1 as a soluble protein exhibits distinct features from the integral membrane protein CAV1 in caveola formation^43^ and signal transduction^18,19^. In this study, our findings suggest that CAVIN1-mediated ROS regulation is independent of CAV1. Specifically, this is because i) CAV1 over-expression did not rescue the reduction of ROS in CAVIN1-null cells (Figure S2B); ii) there is negligible expression of CAV1 paralogs in zebrafish skeletal muscle cells^66,67^, suggesting that the significantly increased ROS scavenging ability observed in *cavin1a/1b* DKO zebrafish skeletal muscle (Figure 2D-E) is a CAV1-independent effect. In addition, CAVIN1 and CAV1 share few similarities in the regulation of NRF2 signaling. Different from our CAVIN1-NRF2 signaling model, a previous study showed that CAV1 is not redistributed and only binds to NRF2 at the PM under oxidative stress^22^. Moreover, CAV1 only affects the subcellular localization but not the half-life of NRF2^22^. This suggests that release from caveolae, interaction with NRF2 in the cytoplasm, and promotion of NRF2 degradation are all unique mechanisms for CAVIN1 in the regulation of NRF2-mediated antioxidant defense.

We demonstrate comprehensively in our cell and zebrafish experiments that the pathophysiological effects associated with ROS dysregulation in cells and tissues lacking CAVIN1 are mechanistically a direct consequence of CAVIN1-NRF2 signalling changes. We show that caveolae directly form a part of a signalling pathway in sensing and mediating acute oxidative stress by releasing CAVIN1 to inactivate the NRF2-dependent antioxidant response, leading to ferroptosis. These findings illustrate a crucial role for caveolae as sensors of cellular stress and demonstrate that under both healthy and disease conditions ROS regulation via the CAVIN1-NRF2 axis is essential for maintaining homeostatic cellular balance by promoting the elimination of severely damaged or malignant cancer cells.

## Acknowledgments

This work was supported by the National Health and Medical Research Council of Australia (grants APP1140064 and APP1150083 and fellowship APP1156489 to R.G.P.; grants APP1125390, APP1140906 to M.T.R. and D.A.S.; fellowship APP1140851 to D.A.S.). R.G.P. is supported by the Australian Research Council (ARC) Centre of Excellence in Convergent Bio-Nano Science and Technology. The authors acknowledge the use of the Microscopy Australia Research Facility at the Center for Microscopy and Microanalysis at The University of Queensland. Confocal microscopy was performed at the Australian Cancer Research Foundation (ACRF)/Institute for Molecular Bioscience (IMB) Dynamic Imaging Facility for Cancer Biology with funding from the ACRF. We thank the Monash FlowCore and the Monash University Biomedical Proteomics Facility for the provision of instrumentation, training, and technical support.

## Author Contributions

Conceptualization, Y.W., Y.-W.L., K.A.M and R.G.P.; Methodology, Y.W., Y.-W.L. and D.A.S.; Formal Analysis, Y.W., Y.-W.L. and D.A.S; Writing – Original Draft, Y.W., Y.-W.L., D.A.S, K.A.M. and R.G.P.; Writing – Review & Editing, Y.W., K.A.M., R.G.P. Y.-W.L., H.P.L., T.E.H, M.T.R. and D.A.S.; Supervision, Y.W. and R.G.P.; Funding Acquisition, R.G.P., M.T.R. and D.A.S.

## Methods

### Zebrafish (Danio rerio)

Zebrafish were raised and maintained according to institutional guidelines (Techniplast recirculating system, 14-h light/ 10-h dark cycle, The University of Queensland). Adults (90 dpf above) were housed in 3 or 8 L tanks with flow at 28.5 °C, late-larval to juvenile stage zebrafish (6 dpf to 45 dpf) were housed in 1 L tanks with flow at 28.5 °C and embryos (up to 5 dpf) were housed in 8 cm Petri dishes in standard E3 media (5 mM NaCl, 0.17 mM KCl, 0.33 mM CaCl_2_ and 0.33 mM MgSO_4_) (Westerfield, 2000) at 28.5 °C (incubated in the dark). All experiments were performed in accordance with relevant guidelines and regulation with the approval of the University of Queensland (UQ) Animal Ethics Committee (Molecular Biosciences Animal Ethics Committee) and UQ Biosafety Committee. The following zebrafish strains were used in this study: wild-type (TAB), an AB/TU line generated by UQ Biological Resources (UQBR) Aquatics, *cavin1b^-/-uq7rp 33^*, *cavin1a^-/-uq10rp^* (this paper) and the *cavin1a/1b* double knockout (DKO) line generated by crossing between the *cavin1a^-/-uq10rp^* and *cavin1b^-/- uq7rp^* lines and incrossing to homozygosity. Only male juvenile and adult zebrafish were used. Zebrafish of developmental stages up to 15 dpf are prior to specific sex determination^68^ and the developmental stages are specifically stated in corresponding figure legends. All zebrafish used in this study were healthy, not involved in previous procedures and drug or test naïve. All zebrafish of the same clutch, and sex or developmental stage were randomly allocated into experimental groups.

### Cell lines

All cancer cell lines were purchased from the American Tissue Culture Collection (ATCC) and grown in recommended medium. Specifically, WT HeLa cells, CAVIN1-null HeLa cells and A431 cells were grown in Dulbecco’s modified Eagle’s medium (DMEM) with 10 % (vol/vol) fetal bovine serum (FBS) at 37 °C in a humidified atmosphere containing 5% CO_2_. MCF-7 cell line was subjected to STR profiling (QIMR Berhofer Medical Research Institute) as described previously^18^. In addition, regular *mycoplasma* test was performed in our laboratory for cell line authentication.

### Animal handling and reagents

Zebrafish embryos up to 7 dpf were raised and handled in standard E3 media during experiments (5 mM NaCl, 0.17 mM KCl, 0.33 mM CaCl2, 0.33 mM MgSO4). All post-embryonic zebrafish measurements were carried out between tanks of similar population densities and conditions. All reagents were obtained from Sigma-Aldrich unless otherwise specified.

### Generation of CAVIN1-null cell lines and cavin1a/1b DKO zebrafish

*CAVIN1*-null cell lines were generated using two independent genomic editing methods. A pair of TALENs were designed to generate a double strand break within human *CAVIN1* gene (see DNA binding sequences of TALENs in Key Sources Table) (Figure S1A-D). The screening strategies have been previously described^69^.

The CRISPR-Cas9 system was used to generate the second *CAVIN1*-null HeLa cell line (Figure S1E-G). Cells at 70% confluency were co-transfected with guide RNA (gRNA, see sequence in Key Sources Table) targeting *CAVIN1* exon 1 region and Cas9 nuclease at a 1:5 ration (μg:μg) using Lipofectamine Cas9 plus reagent. Vector peGFP/NeoR was additionally transfected for the selection of gRNA positive cells. G418 at 2.0 mg/ml was added to the medium for selection on the following day. Replace medium every two days using fresh medium with 2.0 mg/ml G418 until entire untransfected cells were killed. Transfected cells were then diluted and plated into 96 well plate for single clone isolation using array dilution method as described previously^70^. T7 endonuclease I mismatch assays, DNA sequencing assays, TA clone sequencing assays and immunoblotting were performed to validate the editions in *CAVIN1* genes in the cells from different single colony. The clone with homozygous deletion in *CAVIN1* (Δ527-840) was identified and selected for this study.

*Cavin1a/1b* DKO line was generated as previously described^33^. Target site with >50% G/C content and no predicted off-target site for zebrafish *cavin1a* (NCBI reference sequence: XM_001920667.5, corresponding Uniprot accession number: E7F0K3) specific sgRNA was selected using the web tool CHOPCHOP (Montague TG, Cruz JM Gagnon etc CHOPCHOP Nucleic Acids Res 42 2014). Synthesis of sgRNA was carried out in a cloning-independent method as adapted from Gagnon et al^71^ using the cavin1a gene-specific and constant oligos mentioned in the Key Resources Table. Recombinant Cas9 protein containing a nuclear localization signal (PNA Bio Inc) was reconstituted to a solution of 1 mg/mL Cas9 protein in 20 mM HEPES, 150 mM KCl, 1 % sucrose (pH 7.9) and 1 mM DTT. An injection mixture of 700-753 ng/uL Cas9 protein, 200-208 ng/uL sgRNA and 16 % phenol red was prepared and incubated for 5 min at room temperature (RT) to allow for Cas9-sgRNA complex formation. Cavin1a-targeting Cas9-sgRNA was injected into the cytoplasm of the early one-cell stage WT embryos. Injection volumes were calibrated to approximately 600-800 pL of injection mixture per injection.

Founder rate and percentage of mutant allele in f1 progenies was determined via high resolution melt analysis (HRMA). In the DNA preparation step, for whole-embryo tissue collection, selected 48 hpf embryos were anesthetised by rapid cooling and added into the digestion buffer (1 M KCl, 1 M MgCl_2_, 1 M Tris pH 8.3, 10 % NP 40, 10 % Tween-20, 0.1 % gelatine, 20 mg/mL Proteinase K). For juvenile zebrafish tissue collection, selected juvenile zebrafish was anesthetised in ethyl 3-aminobenzoate methanesulfonate (tricaine) solution, before cutting an approximately 3 mm piece of the caudal fin with a sterile razor blade and placing the fin clip in digestion buffer. The mixture was incubated at 60 °C for 1 h before reaction termination at 95 °C for 15 min. Two different HRMA-compatible platforms were used (Applied Biosystems ViiA 7 Real-Time PCR System, using the MeltDoctor HRM Master Mix, and Roche LightCycler 480 System, using the LightCycler 480 High Resolution Melting Master). HRMA primers are indicated in the Key Resources Table. When using the LightCycler® 480 System, the high resolution melt (HRM) step was initiated after a standard PCR amplification step. The HRM step consist of a denaturation step at 95 °C, followed by an annealing step at 65 °C. Melt data acquisition began at 65 °C and ended at 97 °C with 15 fluorescence readings per degree centigrade at a 0.07 °C /s ramp rate. When using the ViiA™ 7 Real-Time PCR System, the HRM step consist of a denaturation step at 95 °C, followed by an annealing step at 60 °C. Melt data acquisition began at 60 °C and ended at 95 °C at a 0.025 °C /s ramp rate. Stable *cavin1a* f1 mutant zebrafish lines were confirmed using Sanger sequencing with *cavin1a* sequencing primers stated in the Key Resources Table. The selected *cavin1a* line [with protein level change *p.(Asp5Leufs*34)*], which was designated as *cavin1a*^-/-*uq*10*rp*^, was then bred to homozygosity. The *cavin1a/1b* DKO line was generated by crossing the *cavin1a^-/-uq10rp^* line to the *cavin1b^-/- uq7rp^* line (Lim et al 2017) and incrossing the offsprings to homozygosity.

### Quantitative mass spectrometry-based comparative proteomic analysis

Samples were prepared for mass spectrometry and analysed as previously described^25^ but with modification for a label free quantification (LFQ) experiment. Briefly, cells were lysed in 1% (w/v) sodium deoxycholate, 100 mM Tris-HCl (pH 8.1), Tris(2-carboxyethy)phosphine (TCEP), 20 mM chloroacetamide and incubated at 99 °C for 10 min. Reduced and alkylated proteins were digested into peptides using trypsin by incubation at 37 °C overnight, according to manufacturer’s instructions (Promega). Sodium deoxycholate was removed from the peptide mixture using SDB-RPS (Merck) stage-tips made in-house as described^25,72^. Peptides were reconstituted in 0.1% trifluoroacetic acid (TFA), 2% ACN and analysed by online nano-HPLC/electrospray ionization-MS/MS on a Q Exactive Plus connected to an Ultimate 3000 HPLC (Thermo-Fisher Scientific). Peptides were loaded onto a trap column (Acclaim C18 PepMap nano Trap x 2 cm, 100 μm I.D, 5 μm particle size and 300 Å pore size; ThermoFisher Scientific) at 15 μL/min for 3 min before switching the pre-column in line with the analytical column (Acclaim RSLC C18 PepMap Acclaim RSLC nanocolumn 75 μm x 50 cm, PepMap100 C18, 3 μm particle size 100 Å pore size; ThermoFisher Scientific). The separation of peptides was performed at 250 nL/min using a non-linear ACN gradient of buffer A (0.1 % FA, 2% ACN) and buffer B (0.1% FA, 80 % ACN), starting at 2.5 % buffer B to 35.4% followed by ramp to 99 % over 278 minutes. Data were collected in positive mode using Data Dependent Acquisition using m/z 375 - 1575 as MS scan range, HCD for MS/MS of the 12 most intense ions with z ≥ 2. Other instrument parameters were: MS1 scan at 70,000 resolution (at 200 m/z), MS maximum injection time 54 ms, AGC target 3E6, Normalized collision energy was at 27% energy, Isolation window of 1.8 Da, MS/MS resolution 17,500, MS/MS AGC target of 2E5, MS/MS maximum injection time 54 ms, minimum intensity was set at 2E3 and dynamic exclusion was set to 15 sec. Raw files were processed using the MaxQuant platform^73^ version 1.6.5.0 using default settings for a label-free experiment with the following changes. The search database was the Uniprot human database containing reviewed canonical sequences (June 2019) and a database containing common contaminants. “Match between runs” was enabled with default settings. Maxquant output (proteinGroups.txt) was processed using Perseus^74^ version 1.6.7.0. Briefly, identifications marked “Only identified by site”, “Reverse”, and “Potential Contaminant” were removed along with identifications made using <2 unique peptides. Log_2_ transformed LFQ Intensity values were grouped into control and knockout groups, each consisting of three replicates. Proteins not quantified in at least two replicates from each group were removed from the analysis. Annotations (Gene Ontology (GO), Biological Process (BP), Molecular Function (MF), Cellular Compartment (CC), KEGG, Corum and PFAM) were loaded through matching database built into Perseus with the majority protein ID. Pathway analysis (Table S2; Figure 1B) and upstream regulator analysis (URA) (Table 1; Figure 1C) were performed on significantly altered proteins in CAVIN1-null cells using the “core analysis” function included in the Ingenuity Pathway Analysis (IPA) software (QIAGEN Bioinformatics, content version 44691306)^75^. Overlap *p*-value in URA measures the significance between the dataset genes and the genes that are regulated by a transcriptional regulator. It is calculated using Fisher’s Exact Test and significance is generally attributed to *p*-values<0.01. Enrichment analysis (Figure 1D-E) was performed using online tool “inBio Discover™”.

### SDS PAGE and western blot analysis

Cell lysates were determined using a BCA Protein Assay Kit as the standard. Estimated proteins (20-40 μg) were separated by SDS PAGE and transferred to PVDF membranes. Western blots were performed as standard procedures. Detection and quantification of target proteins was carried out on a scanning system (BIO-RAD, ChemiDoc^TM^ MP) using horseradish peroxidase (HRP)-conjugated secondary antibodies and ECL detection reagent. Intensities of bands were quantitated by ImageJ 2.0 software.

### Proximity ligation assay

Duolink II Detection Kit was utilized to assess protein-protein proximity according to the manufacturer’s instruction. Primary antibodies with different species were used for the detection of each protein pair. Specifically, Rabbit anti-NRF2 (Abcam) and mouse anti-Ubiquitin (Sapphire Bioscience) for PLA in Figure 3I and 3J; Rabbit anti-CAVIN1(rabbit, Proteintech) and mouse anti-NRF2 (Abcam) for PLA in Figure 4J and 4T. PLA signals were visualized and imaged by Zeiss LSM 880 confocal microscope and quantified using ImageJ 2.0 software.

### Immunofluorescence

Cells were plated onto glass coverslips at 70 % confluence and then fixed in 4 % (vol/vol) paraformaldehyde in PBS for 15 min at RT after the indicated treatments. Coverslips were washed three times in PBS and permeabilized in 0.1 % (vol/vol) Triton X-100 in PBS for 5 min and blocked in 1 % (vol/vol) BSA in PBS for 30 min at RT. Primary antibodies were diluted in 1% BSA/PBS solution at optimized concentrations and incubated with coverslips for at 4°C overnight. Diluted secondary antibodies (1:500 in 1% BSA/PBS) conjugated with fluorescent dyes were later added onto coverslips and incubated for 1 h. Coverslips were washed in PBS for three times and mounted in Mowiol in 0.2 M Tris-HCl pH 8.5. The images were taken on Zeiss LSM 880 confocal microscope and intensities of fluorescence were quantitated using ImageJ 2.0.

### Co-immunoprecipitation assay

HeLa cells were seeded in 10-cm dishes (1 x 10^6^ cells per dish) and lysed in ice-cold lysis buffer (20 mM Tris-HCl, pH 7.4, 150 mM NaCl, 10% Glycerol, 1% Triton X-100, cOmplete^TM^ protease inhibitor cocktail tablet from Roche and a PhosSTOP tablet from Roche). Lysates containing 500-1000 μg protein (made up to 500 μl by lysis buffer) was pre-cleared using protein A-coupled sepharose beads (20 ul). After centrifuging at 2,500 g for 1 min, supernatants were collected and mixed with 1.5 μg rabbit anti-NRF2 antibody (Abcam) or rabbit anti-CAVIN1 antibody (Abcam). Rabbit anti-IgG (1.5 μg/500 μl) was used as negative control. Incubate the lysates with antibodies over night at 4 °C and protein A (20 μl) was added for another 3 h incubation. The tubes were centrifuged and the supernatant was removed from the beads. After the beads were washed with lysis buffer for three times, sample buffer was added to denature the samples by boiling it at 95 °C for 10 min. Western blot analysis was applied to detect the proteins that co-immunoprecipitated with NRF2 or CAVIN1.

### RNA preparation, reverse-transcription and quantitative real-time PCR

For human cell lines, total RNA was extracted using QIAGEN RNA isolation kit following the manufacturer’s instructions and measured using a NanoDrop spectrophotometer (ThermoFisher Scientific). Two-step reverse transcription was conducted to access single strand cDNA using SuperscriptIII reverse transcriptase (ThermoFisher Scientific) as per manufacturer’s instruction. SYBR-Green PCR Mater Mix or Taqman (CAVIN1 probe) was utilized for real-time PCR using a ViiA7 Real-time PCR system (ThermoFisher Scientific). The sequence of oligonucleotide primers for real-time PCR are listed in Key Resources Table. Gene expression was analysed using the delta-delta Ct method as previously described^76^.

For zebrafish, RNA was isolated from *cavin1a/1b* DKO and WT zebrafish embryos (> 100 embryos randomly selected from 1 clutch) using the RNeasy Mini Kit (QIAGEN) and cDNA synthesis was performed using SuperscriptIII reverse transcriptase (ThermoFisher Scientific) as per manufacturer’s instructions. Quantitative real-time PCR was performed using the SYBR Green PCR Master Mix on a ViiA7 Real-time PCR system (ThermoFisher Scientific) according to manufacturer’s instructions with 3 biological replicates (embryos randomly selected from 3 clutches) and 3 technical replicates on 96-well plates. Primers used were detailed in the Key Resources Table. Gene expression was analyzed using the delta-delta Ct method as previously described^76^.

### Live imaging of zebrafish embryos

Prior to imaging, zebrafish (up to 26 dpf) were anesthetized in tricaine solution in E3, mounted in 1% low melting point (LMP) agarose on MatTek glass bottom dishes in a lateral view unless otherwise stated (anterior to the left, posterior to the right) and submerged in tricaine solution. The embryos were submerged in tricaine solution in E3 throughout all imaging periods.

For stereo microscopy of general morphology, zebrafish embryos were mounted as described above and imaged under a SMZ1500 stereomicroscope. For fluorescence stereo microscopy, zebrafish embryos were mounted as described above and imaged under a Nikon SMZ18 stereo microscope.

For live confocal images of WT and *cavin1a/1b* DKO zebrafish, embryos were incubated at 28°C and mounted in LMP agarose as above and imaged under a Zeiss LSM880 confocal microscope. For general characterization of zebrafish, WT and *cavin1a/1b* DKO embryos were preincubated in BODIPY FL C5-Ceramide (ThermoFisher Scientific) for 24 h (at 28 °C) s previously described^77^ before the anaesthesia and mounting steps detailed above.

### Body length measurement of live zebrafish

Images captured using the NIS Elements Version 4.20 software on the SMZ1500 stereomicroscope were used to measure the body length of 3 dpf WT and *cavin1a/1b* DKO zebrafish embryos. Body length was defined as the region from the tip of the anterior end of the embryo to the end of the trunk before the caudal fin. Measurements were conducted using Fiji and were non-blind. Embryos were randomly selected from 3 biological replicates (clutches) with no prior formal sample-size estimation.

### Reactive oxygen species (ROS) detection

The cell-permeant reagent H_2_DCFDA (2’, 7’-dichlorodihydrofluorescein diacetate) (Thermo Fisher Scientific) was employed to represent the ROS levels in HeLa cells. Cells were lysed after incubation with reagent (20 μM) for 30 min at 37°C. Cell lysates of untreated or H_2_O_2_ treated HeLa cells containing equal protein (100 μg) were diluted to 15 μl working volume and loaded into a flat-bottom white 384 well plate, which was then applied to a highly sensitive top-read microplate reader (TECAN) that does not cause photo-oxidation effect on H_2_DCFDA probes^78^ for the quantification of fluorescence (Excitation/Emission in nm: 485/528) at a single time point. All the samples were kept in the dark prior to measurement. Fluorescence values for each group were corrected by subtracting background fluorescence value generated by a cell lysis buffer alone control. ROS were also labelled by CellROX Green probe (Thermo Fisher Scientific) and visualized as green fluorescence in mounted cultures by Zeiss LSM 880 confocal microscope on 63x oil objective lens.

### Lipid peroxidation detection

LPO in cells with were detected using Image-iT lipid peroxidation kit (ThermoFisher) according to the manufacturer’s instruction. For the inhibition of LPO, cells were pre-treated with α-Tocopherol for 30 min at 25 μM as this concentration of α-Tocopherol most effectively protected lipids from oxidant attack (Figure S4F). BODIPY-C11 fluorescence was visualised in live cells by Zeiss LSM 880 confocal microscope on 63x oil objective lens.

### ROS assessment in zebrafish embryos

2 dpf zebrafish were incubated in 2 mM H_2_O_2_ for 1 h, washed in E3 media 3 times and stained using H_2_DCFDA (20 µM in E3 media) for 30 min. The embryos were then washed with E3 media 2 times, anesthetized in tricaine and mounted in 1% LMP agarose on 35 mm MatTek glass bottom dishes for imaging under a Zeiss LSM880 confocal microscope or Nikon SMZ18 stereo microscope. To quantitate, stereo microscope images of treated and untreated zebrafish were analysed using Fiji. The lookup table of the images were inverted for visualization of embryo somite segments. For each embryo, a constant area of 4 somite pairs at the end of the embryo yolk extension was analyzed and the corrected total fluorescence (integrated density - [area of selection * mean grey value of image background]) was calculated. DCF fluorescence of individual H_2_O_2_-treated zebrafish was normalized using the mean corrected total fluorescence of its corresponding untreated clutch.

### Tail excision in zebrafish embryos

2 dpf zebrafish were anesthetized in tricaine and placed on an open petri dish in a drop of tricaine solution in E3. Tail amputation was carried out using a scalpel to excise the tail end of the embryo at a constant region of three to six rows of notochord cells away from the posterior tip of the notochord.

For regeneration assay, regrowth area was measured at indicated time points post tail excision and then divided by the tail area before excision. Regeneration efficiency is expressed as a percentage (%) of the tail area before excision.

For ROS measurement, embryos were treated 30 min before imaging with H_2_DCFDA (20 µM in E3 media) as above. To quantitate, integrated DCF intensity in a constant area at the wound site, as well as in the normal tissue, was analyzed using Image J. ROS accumulation following injury is indicated by the value of DCF intensity in the injured tissue that was normalized with the DCF intensity in the normal tissue within the same zebrafish.

### TUNEL assay in zebrafish embryos

Zebrafish at 4hpe was fixed with 100% methanol and stored at -20°C overnight. On the following day, methanol was gradually removed by washing zebrafish with 70%, 50%, 25% methanol and PBS three times. Apoptotic cells in zebrafish were then detected using TUNEL assay kit (*In Situ* Cell Death Detection Kit, TMR red) according to the manufacturer’s instruction. Reagents required for TUNEL assay was prepared freshly before each experiment. Fixed embryos were then processed as described above for confocal imaging under a Zeiss LSM880 confocal microscope.

### Hydrogen peroxide scavenging assay

Cell lysate containing 2 mg protein in 1.5 mL distilled H_2_O were incubated with 62.5 μl H_2_O_2_ (5 mM) at 37°C for 15 min to allow redox reaction to occur. Next, 0.25 mL ammonium iron (II) sulphate hexahydrate (1 mM) was added to reaction solution for another 5 min incubation. 1.5 ml 1,10-phenanthroline monohydrate (1 mM) were added at last and the amount of tri-phenanthroline complex that formed by ferrous and phenanthroline were indicated by optical density value that measured at 510 nm. hydrogen peroxide scavenging activity was calculated using following formula:

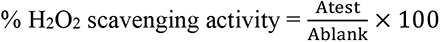

A_blank_ is the absorbance of solution containing only ferrous ammonium sulphate and 1,10-phenanthroline.

### Cell viability assay

Cells were counted using a hemocytometer and seeded into 96-well plate at 5000 cells in 90 μl medium per well. After 6, 12, 24, 48 and 72 h incubation, 10 μl of PrestoBlue™ Viability Reagent (10x) (absorbance wavelength: 600 nm) (Thermo Fisher Scientific Inc.), which was quickly converted by viable cells to a red fluorescence reduced form of the dye possessing an absorbance at 570 nm, was added into each well and incubated for 30 min. Both absorbance value at 570 nm and 600 nm were measured for each plate, where 570 nm was used as experimental wavelength and 600 nm as normalization wavelength. The absorbance values for wells only containing medium without cells were read for background correction. Raw data was processed to evaluate the percent reduction of PrestoBlue™ reagent for each well by using the following equation referring to the manufacturer’s protocol:

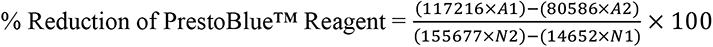

Where: A1 = Absorbance of test wells at 570 nm, A2 = Absorbance of test wells at 600 nm, N1 = Absorbance of media only wells at 570 nm, N2 = Absorbance of media only wells at 600 nm.

### Live-cell imaging and laser treatment

HeLa cells were plated to 35-mm glass bottom dishes the day before imaging. CO_2_-independent medium (Gibco) containing 10% FBS was applied on cells to remain the cell viability. Cells were incubated with C11-BODIPY probes for 30 min for lipid labelling prior to imaging. Zeiss LSM 880 confocal microscope with Airyscan was utilised for imaging. After 405 nm laser pulses applied to cells, FITC and Texas Red channels were used to collect images over 80 sec with 1 sec interval time. Image analysis was performed using Image J 2.0. ROIs were done and imported from Zeiss Zen 2012 Black to locate laser-treated region.

### Nile red vital staining of zebrafish juveniles

Live 26 dpf WT and *cavin1/a1b* DKO juvenile zebrafish were incubated in system water supplemented with Nile Red stock (1.5 mg/mL Nile Red in acetone) to a working concentration of 0.4 μg/mL in the dark for 35 min. Zebrafish were then anesthetized in tricaine, washed for 1 min in tricaine solution, mounted in 3% LMP agarose on a 35 mm petri dish and imaged under a Zeiss LSM 510 Meta inverted confocal microscope.

### Electron microscopy of adult zebrafish

This protocol is a modified version originally by Deerinck et al. (2010) and designed to enhance membrane contrast using reduced osmium tetroxide, thiocarbohydrazide-osmium, uranyl acetate and en bloc lead nitrate staining. A solution containing 2.5 % glutaraldehyde in 2X PBS was added to the dish in equal volume with dissected adult zebrafish tissues and placed for 5 min in a Pelco Biowave under vacuum and irradiated at 80 W. Dissected adult tissues were then irradiated in fresh fixative (2.5 % glutaraldehyde), under vacuum, for a further 6 min. Embryos or adult tissues were washed 4 x 2 min in 0.1 M cacodylate buffer. A solution containing both potassium ferricyanide (1.5%) and osmium tetroxide (2 %) in 0.1 M cacodylate buffer was prepared and samples immersed for 30 min at RT. Following 6 x 3 min washes in distilled water, samples were then incubated in a filtered solution containing thiocarbohydrazide (1%) for 30 min at RT. After subsequent washing in distilled water (6 x 2 min), samples were incubated in an aqueous solution of osmium tetroxide (2 %) for 30 min. Samples were washed again in distilled water (6 x 2 min) and incubated in 1% uranyl acetate (aqueous) for 30 min at 4^°^C. Further distilled water washes (2 x 2 min) were completed before adding a freshly prepared filtered 0.06% lead nitrate in aspartic acid (pH 5.5) solution warmed to 60°C. The lead nitrate solution containing tissue blocks was further incubated for 20 min at 60°C before rinsing in distilled water (6 x 3 min) at RT. Samples were dehydrated twice in each ethanol solution of 30%, 50%, 70%, 90% and 100% absolute ethanol for 40 s at 250 watt in the Pelco Biowave. Epon LX112 resin was used for embedding the tissue with infiltration steps at 25%, 50%, 75% resin to ethanol in the Pelco Biowave under vacuum at 250 watt for 3 min and finishing with 100% resin (twice), before the final resin embedding and placed in a 60 oC oven for 12 hours. Blocks were sectioned on a Leica UC64 ultramicrotome at 60 nm and mounted on formvar coated 3 slot Cu grids. Thin sections (60 nm) were viewed on a Jeol JEM-1011 at 80kV.

### Quantification and Statistical Analysis

GraphPad Prism software version 9.0 was used for statistical analysis. An unpaired two-tailed Student’s *t*-test was used to determine significance between two groups. Significance in multiple groups was assessed by one-way or two-way analysis of variance (ANOVA), followed by Tukey’s, Dunnett’s or Sidak’s test, as specified in the legends. Quantified values are presented as mean ± SD from at least n = 3 biological replicates. All tests used a significance (α) level of 0.05. The exact *p*-values are reported in the figures. If the *p*-value is less than 0.0001, we report “****” in the figures. All of the statistical details of experiments can be found in the figure legends.

### Data availability

All data supporting the findings of this study are available from the corresponding author on reasonable request. Comparative proteomics dataset generated during this study will be upload to PRIDE upon publication. Raw data of western blot analysis from Figures 2, 3, 4, 5 and S1 deposited on figshare repository can be previewed at: [https://figshare.com/s/c70e582f1dec5bfc42cb].

### Code availability

This study did not use any unreported custom code or mathematical algorithm that is deemed central to the conclusion.

## Key Resource Table

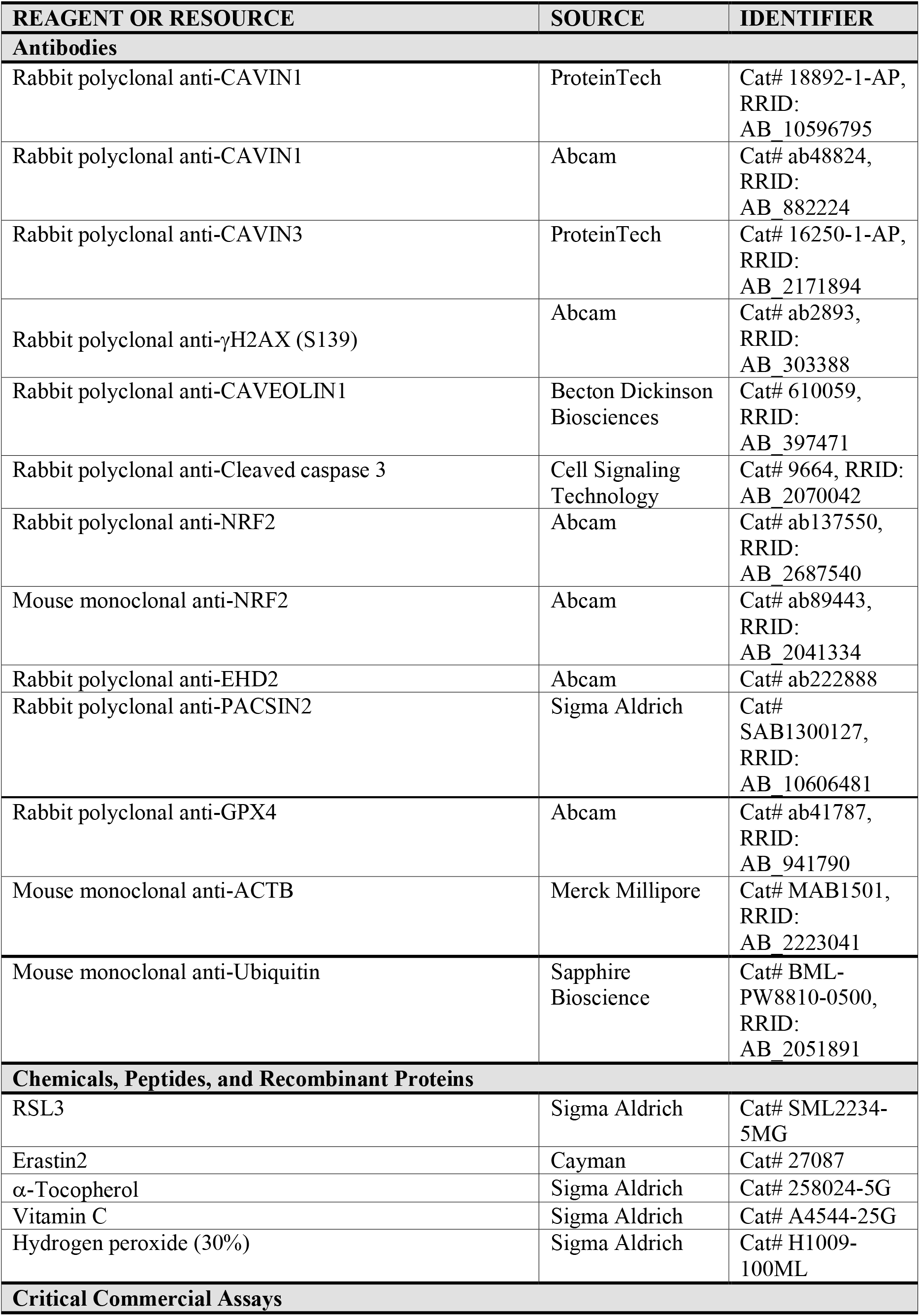
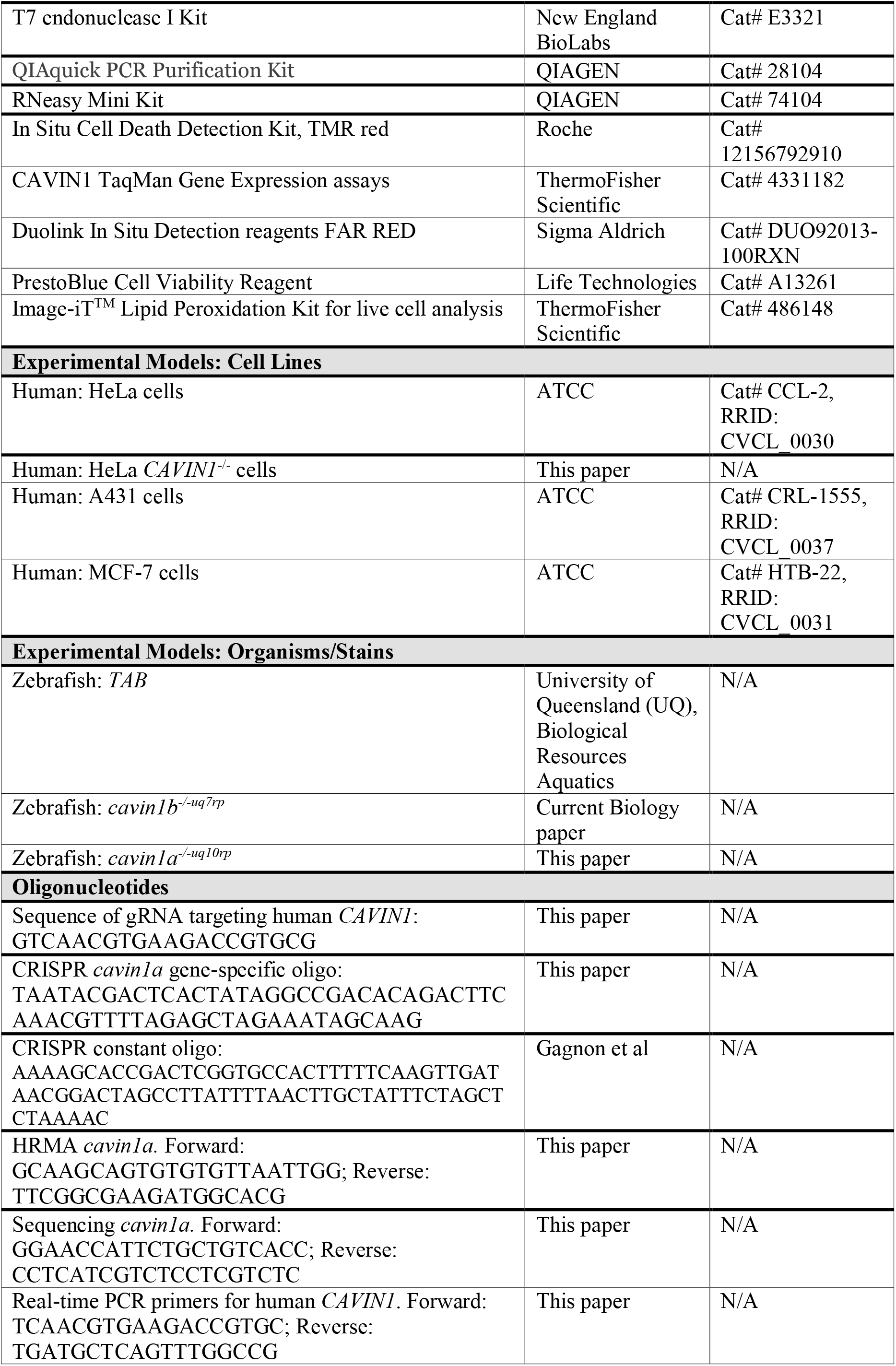
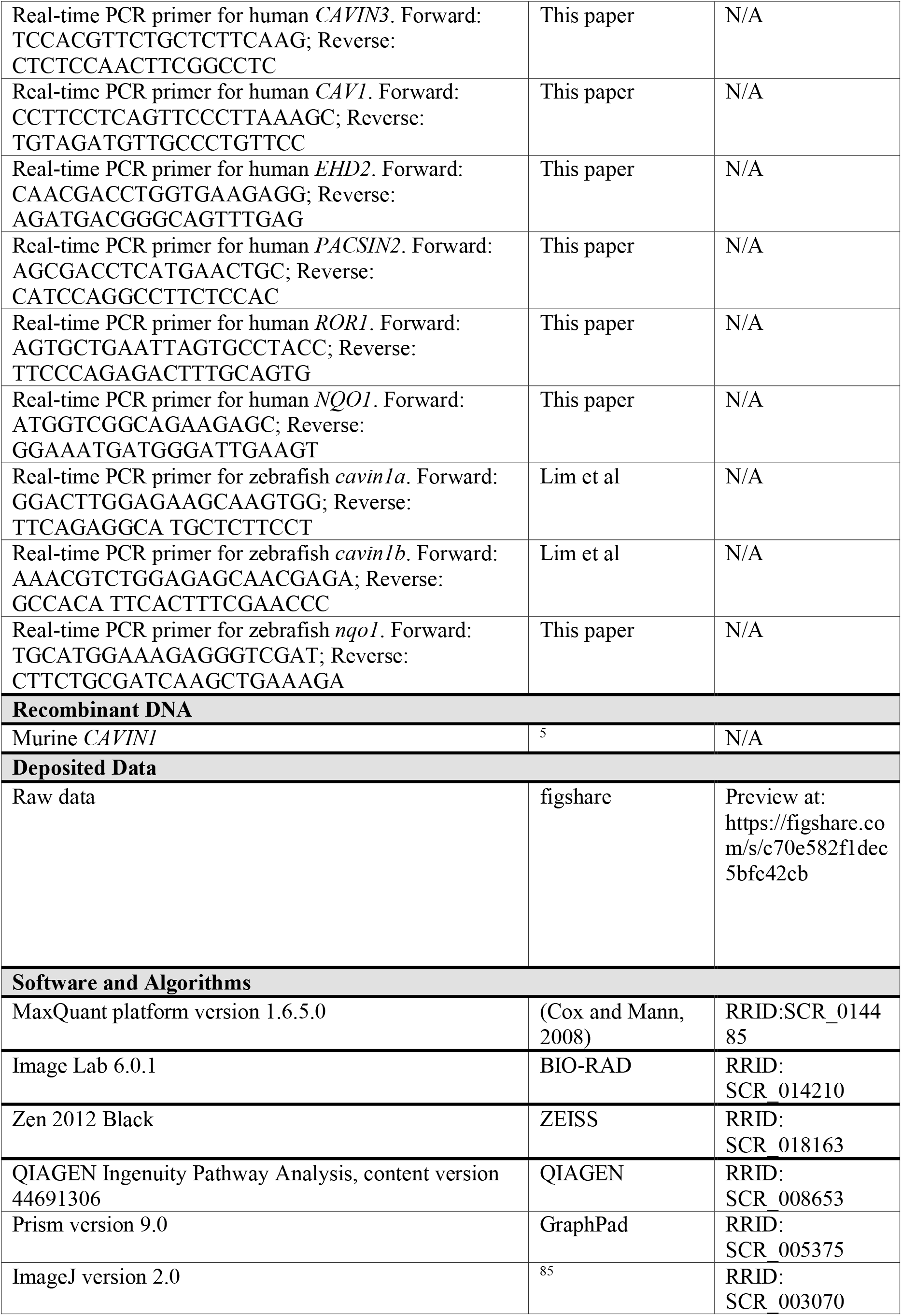

