## Supplementary Information for "Caveolae respond to acute oxidative stress through membrane lipid peroxidation, cytosolic release of CAVIN1, and downstream regulation of NRF2"

This PDF file includes:

- Supplementary Figures 1 to 6
- Titles and Legends for Supplementary Tables 1 to 2
- Supplementary Text

Other Supplementary Information for this manuscript include the following:

- Supplementary Tables 1 to 2
- Supplementary Videos 1 to 2

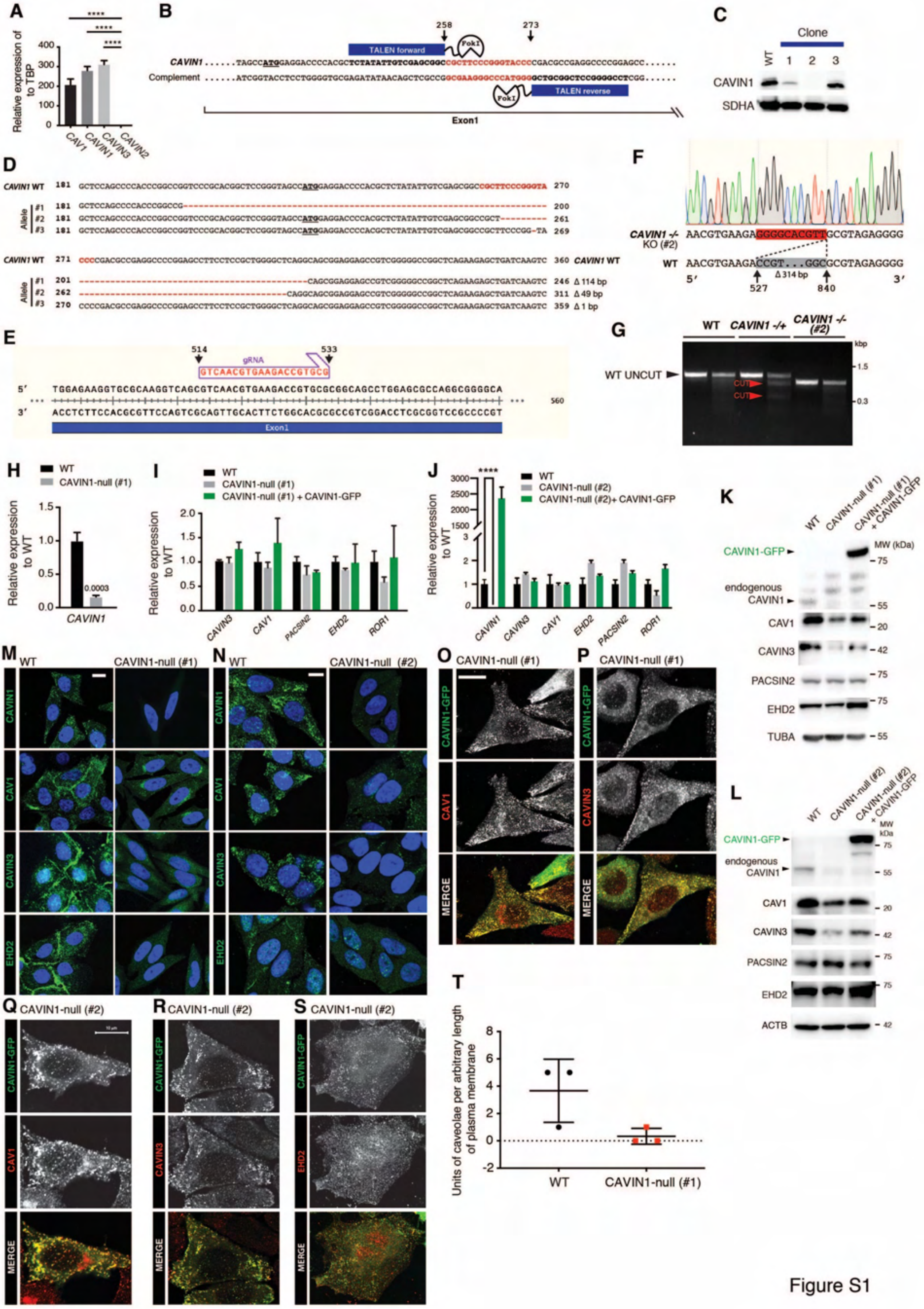

Figure S1

**Figure S1.** Generation and characterization of CAVIN1-null cell lines. (A) Transcript levels of *CAV1*, *CAVIN1*, *CAVIN2* and *CAVIN3* in HeLa cells assessed by real-time PCR. (B) TALEN-targeting region is highlighted in red in human *CAVIN1* human gene sequence. (C) Western blot detection of CAVIN1 protein in different clones. (D) Deletions (dashes) detected in a total of six bacterial clones by DNA sequencing. The region marked in red indicates the TALEN-targeting sites. (E) Designed guide RNA sequence and target sites in human *CAVIN1* gene. (F) DNA sequencing revealing a deletion in *CAVIN1* gene in selected clone. (G) T7 endonuclease mismatch assays were performed to evaluate the gene editing efficiency of CRISPR/Cas9 in selected clones. (H-I) The mRNA levels of *CAVIN1* (Taqman) (H) and caveola associated components *CAVIN3*, *CAV1*, *EHD2*, *PACSIN2* and *ROR1* (SYBR Green) (I) were evaluated in WT, CAVIN1-null (#1) and CAVIN1-GFP expressing CAVIN1-null (#1) cells by real-time PCR assays. Statistical significance was determined by Student's *t*-test and two-way ANOVA for (H) and (I) respectively. (J) The mRNA levels of *CAVIN1* and caveola associated components were evaluated in WT, CAVIN1-null (#2) and CAVIN1-GFP expressing CAVIN1-null (#2) cells by real-time PCR assays (SYBR Green) and compared using two-way ANOVA. (K-L) The protein levels of CAVIN1 and caveola associated components were analyzed in WT, CAVIN1-null and CAVIN1-GFP expressing CAVIN1-null cells by western blotting, n=3 independent experiments. (M-N) Immunofluorescence showing the localization of CAVIN1, CAV1, CAVIN3 and EHD2 in HeLa WT and CAVIN1-null cells. DAPI (blue) was stained to indicate the nucleus. Scale bar, 10 µm. (O-P) Immunuofluorescence detected the localization of CAV1 (O) and CAVIN3 (P) in CAVIN1-GFP re-expressed CAVIN1-null (#1) cell line. Scale bar, 10 µm. (Q-S) Subcellular localization of CAV1, CAVIN3 and EHD2 in CAVIN1-GFP re-expressed CAVIN1-null (#2) cells was assessed by immunofluorescence. Scale bar, 10 µm. (T) Quantification of the number of caveolae per cell from EM images.

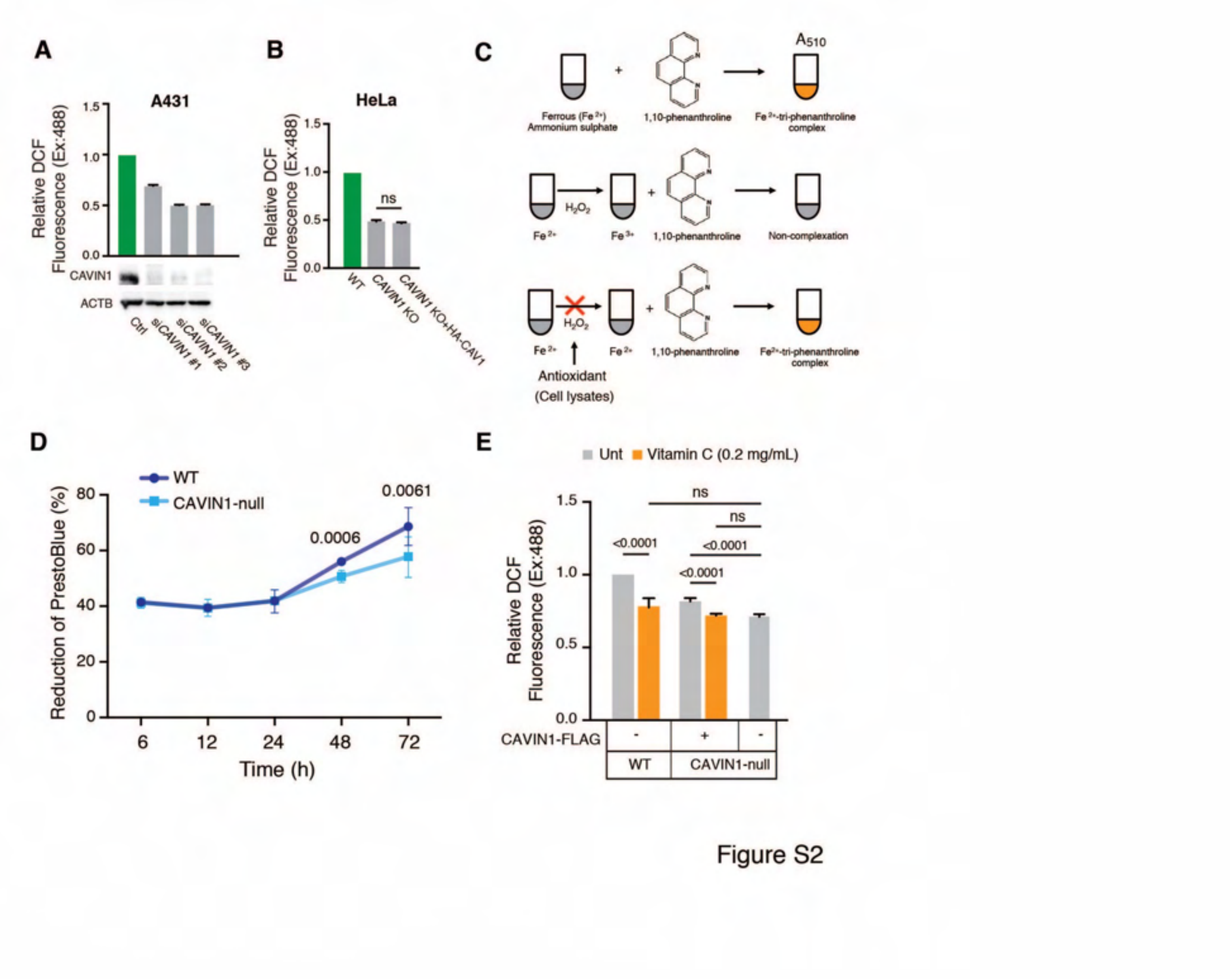


**Figure 2.** Assessment of the roles of CAVIN1 in ROS regulation. (A) DCF fluorescence in A431 cells with or without the depletion of CAVIN1. Three sets of siRNAs were used for comparison. (B) Effect of re-expression of CAV1 on ROS levels in CAVIN1-null cells was indicated by DCF fluorescence intensities. (C) A graphical depiction of the $H_2O_2$ scavenging assay (D) PrestoBlue assays show cell viability (5,000 cells per well for a 96-well plate). Statistical difference between WT and CAVIN1-null cells at each time point was analyzed using two-way ANOVA. (E) Relative DCF fluorescence intensity was calculated and compared using two-way ANOVA, ns=no significance.

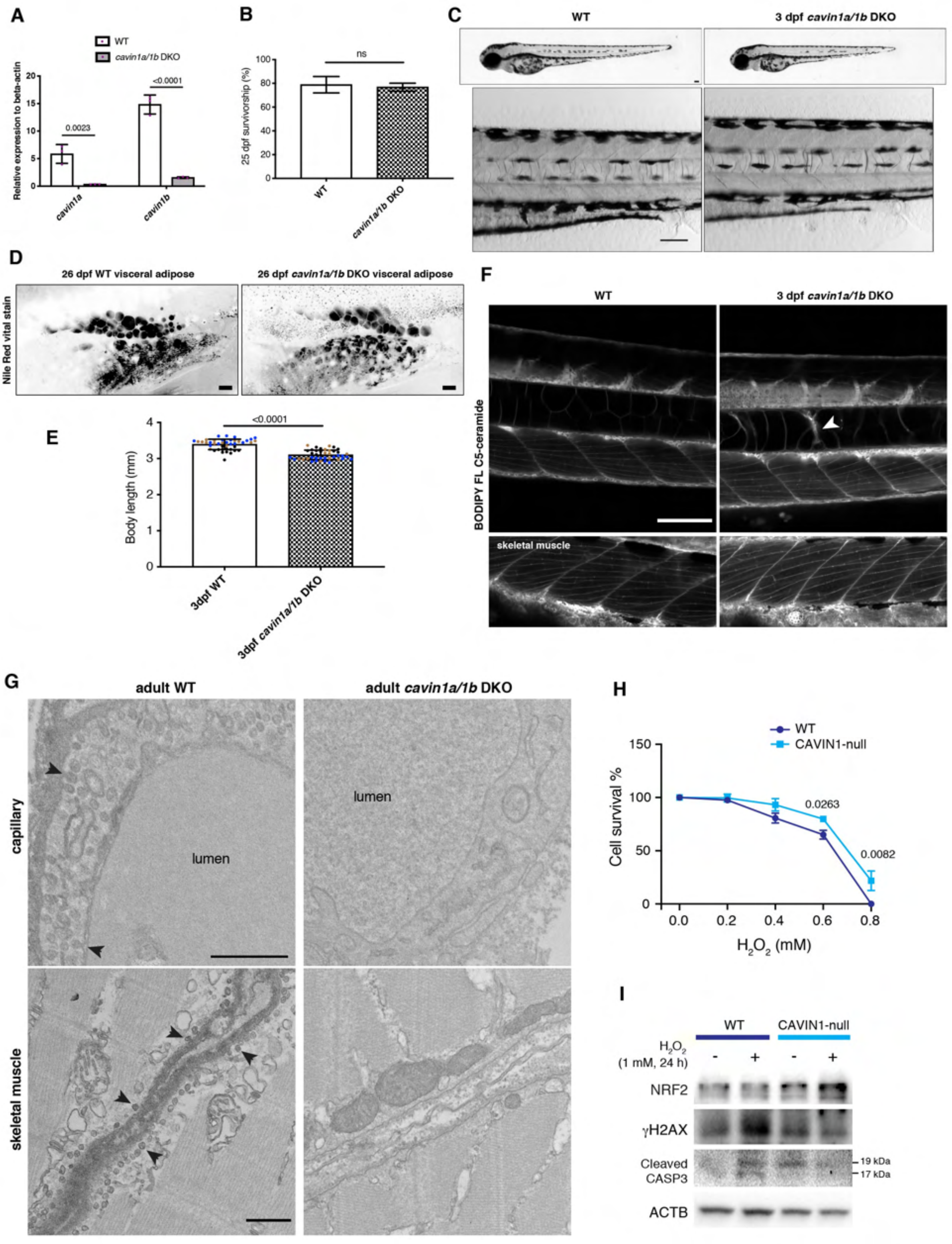

A
WT
cavin1a/1b DKO
Relative expression to beta-actin
0.0023
<0.0001
cavin1a
cavin1b
B
ns
25 dpf survivorship (%)
WT
cavin1a/1b DKO
C
WT
3 dpf cavin1a/1b DKO
D
26 dpf WT visceral adipose
26 dpf cavin1a/1b DKO visceral adipose
Nile Red vital stain
E
<0.0001
Body length (mm)
3dpf WT
3dpf cavin1a/1b DKO
F
WT
3 dpf cavin1a/1b DKO
BODIPY FL C5-ceramide
skeletal muscle
G
adult WT
adult cavin1a/1b DKO
capillary
lumen
lumen
skeletal muscle
H
WT
CAVIN1-null
Cell survival %
0.0263
0.0082
H2O2 (mM)
I
WT
CAVIN1-null
H2O2 (1 mM, 24 h)
NRF2
γH2AX
Cleaved CASP3
19 kDa
17 kDa
ACTB


Figure S3

Despite a significant downregulation of both *cavin1* paralogs indicating nonsense-mediated decay[79], the *cavin1a/1b* DKO line was viable, with no observable defects in early survivorship, gross morphology or adipose development (Figure S3A-D), similar to our single *cavin1b* mutants[33]. Also similar to the single *cavin1b* mutants, *cavin1a/1b* DKO embryos possess defects in body length and notochord lesions (Figure S3E-F).

**Figure S3.** Generation and characterization of *cavin1a/1b* DKO zebrafish. (A) mRNA expression levels of *cavin1a* and *cavin1b* in 5 dpf WT and *cavin1a/1b* DKO zebrafish (relative to β-actin). n = 3 clutches, performed in triplicate. Two-way ANOVA was used for statistical analysis. (B) Survivability of *cavin1a/1b* DKO zebrafish grown to 25 dpf compared to WT (n=71 zebrafish [WT] and n=69 zebrafish [*cavin1a/1b* DKO]; clutch number for both lines=3). (C) Gross morphology of live WT and *cavin1a/1b* DKO zebrafish at 3 dpf. Scale bar, 100 μm. (D) Representative images of Nile Red staining of 26 dpf live WT and *cavin1a/1b* DKO zebrafish visceral adipose. Scale bar, 100 μm. (E) Body length (mm) of 3 dpf WT and *cavin1a/1b* DKO zebrafish (n=42 zebrafish [WT] and n=42 zebrafish [*cavin1a/1b* DKO]; clutch number for both lines = 3, colored dots represent different clutches). (F) Representative live confocal images of BODIPY FL C5-ceramide labelled WT and *cavin1a/1b* DKO zebrafish. Medial view of the zebrafish notochord and skeletal muscles. Arrowhead indicates a notochord lesion. Scale bar, 100 μm. (G) TEM micrographs of skeletal muscle capillary and skeletal muscle of adult male WT and *cavin1a/1b* DKO zebrafish (approximately 3 months post fertilization). Arrowhead indicates caveola. Capillary scale bar, 500 nm; Skeletal muscle scale bar, 1 μm. (H) PrestoBlue assays assessed cell viability after exposure to $H_2O_2$ for 24 h. Relative values compared to untreated HeLa cells are represented as the survival curve (%). Two-way ANOVA was used to compare the values at each time point. (I) NRF2, γH2AX and cleaved CASP3 (caspase 3) protein levels in HeLa cells were assessed by western blot assays. ACTB (β-actin) was detected as loading control.

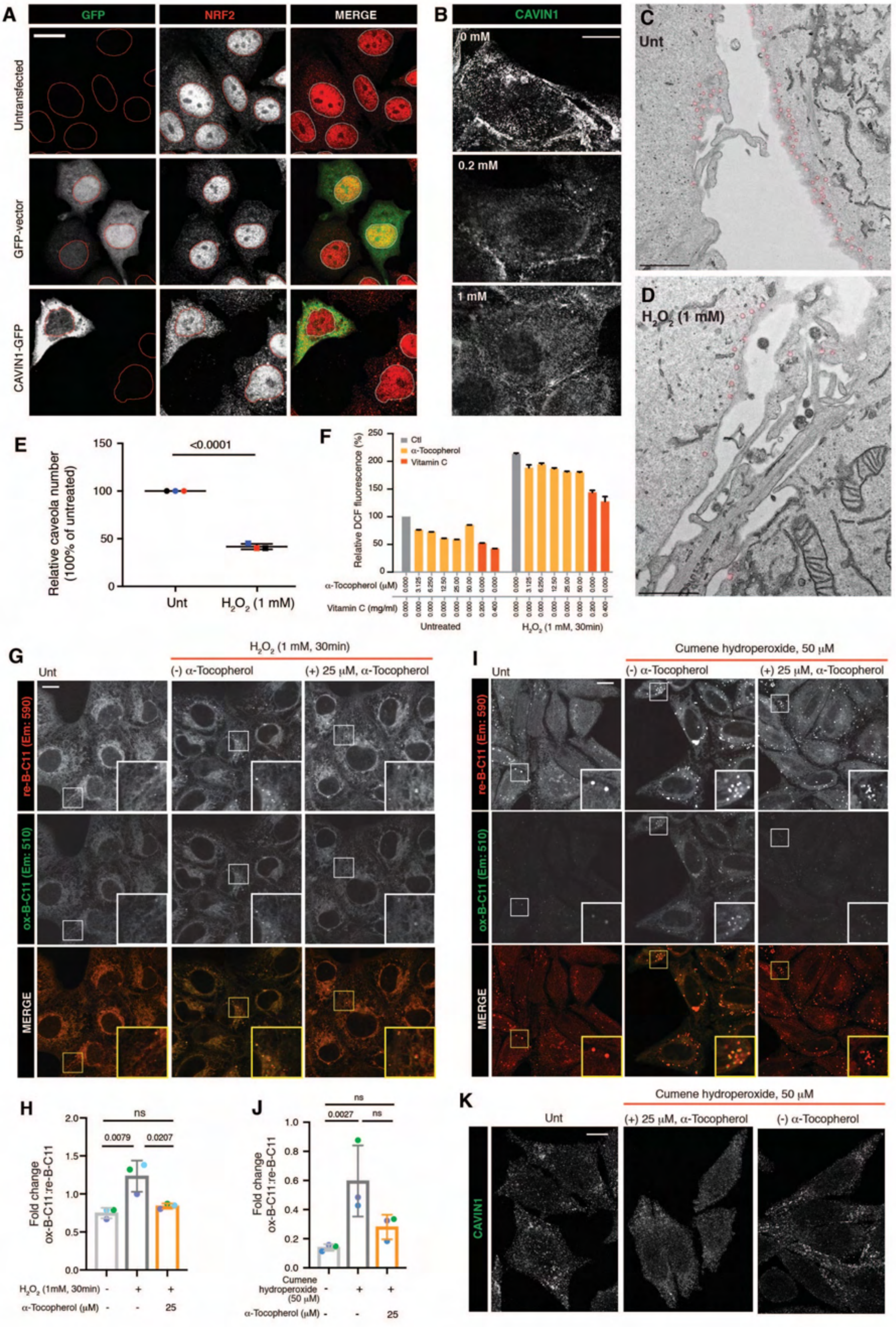
A
GFP
NRF2
MERGE
Untransfected
GFP-vector
CAVIN1-GFP
B
CAVIN1
0 mM
0.2 mM
1 mM
C
Unt
D
H2O2 (1 mM)
E
Relative caveola number (100% of untreated)
<0.0001
Unt
H2O2 (1 mM)
F
Relative DCF fluorescence (%)
Ctl
α-Tocopherol
Vitamin C
α-Tocopherol (μM)
Vitamin C (mg/ml)
Untreated
H2O2 (1 mM, 30min)
G
H2O2 (1 mM, 30min)
Unt
(-) α-Tocopherol
(+) 25 μM, α-Tocopherol
re-B-C11 (Em: 590)
ox-B-C11 (Em: 510)
MERGE
I
Cumene hydroperoxide, 50 μM
H
Fold change ox-B-C11:re-B-C11
ns
0.0079
0.0207
H2O2 (1mM, 30min)
α-Tocopherol (μM)
J
0.0027
Cumene hydroperoxide (50 μM)
K
CAVIN1

Figure S4

**Figure S4.** Lipid peroxidation is responsible for CAVIN1 release and allows for its interaction with NRF2 upon oxidative stress. (A) NRF2 localization in CAVIN1-deficient MCF-7 cells with or without re-expression of CAVIN1-GFP. Scale bar, 10 μm. (B) Representative confocal images of immunofluorescence showing CAVIN1 distribution before and after treatment of $H_2O_2$. Scale bar, 10 μm. (C-D) EM images displaying caveolar levels in untreated and $H_2O_2$ treated A431 cells. Caveolae are highlighted in red. (E) Quantification of caveola number in A431 cells from EM images was analyzed using Student's *t*-test. (F) Effects of α-Tocopherol at different concentrations on the inhibition of ROS levels in HeLa cells. Vitamin C treatment was included as a positive control. (G) B-C11 signal was observed using confocal microscopy in A431 cells with different treatments. Scale bar, 10 μm. (H) Ratio values (ox-B-C11:re-B-C11) for each group in (G) were statistically analyzed by using two-way ANOVA. (I) Confocal images showing reduced and oxidized signals of B-C11 probes in HeLa cells with different treatments. Cumene hydroperoxide (50 μM) was used for lipid peroxidation induction. Scale bar, 10 μm. (J) The (ox-B-C11:re-B-C11) ratio for each group in (I) was compared using one-way ANOVA. (K) Immunofluorescence showing CAVIN1 distribution following cumene hydroperoxide treatment in HeLa WT cells with or without pretreatment by α-Tocopherol. Scale bar, 10 μm.

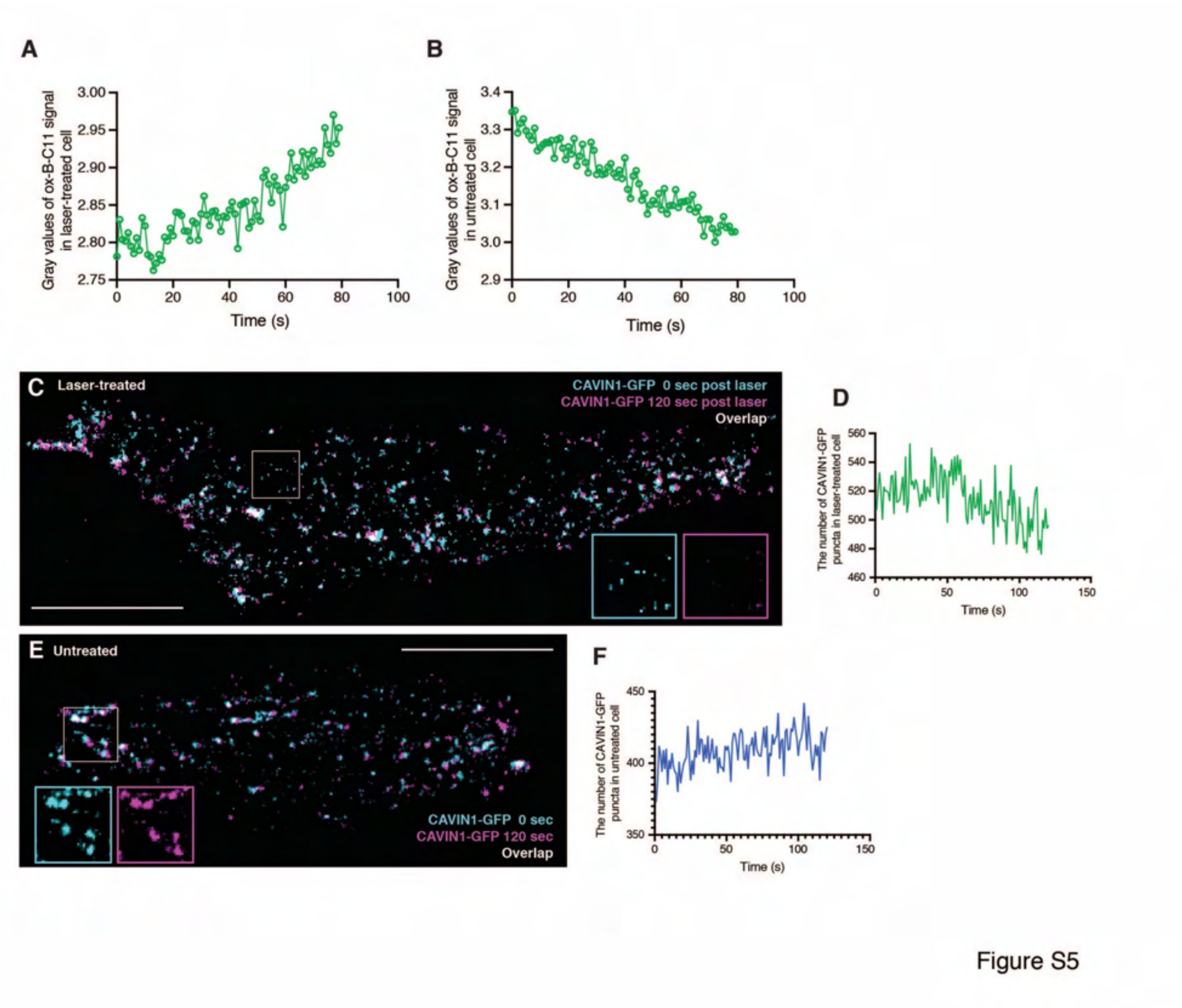


**Figure S5.** Lipid peroxidation leads to CAVIN1 dissociation from caveolae. (A-B) Gray values of ox-B-C11 over 2 min in laser-treated HeLa cell (A, see movie in Video S1) and in untreated HeLa cell (B, see movie in Video S2). (C) Merged images obtained at 0 sec (cyan) and 120 sec (meganta) following laser treatment showing the overlap of CAVIN1-GFP puncta at the PM, see full movie in Video S3. Scale bar, 10 µm. (D) The changes in the number of CAVIN-GFP puncta in laser-treated cells over 120 sec following laser treatment. (E) Overlapped images acquired at 0 sec (cyan) and 120 sec (meganta) during live-cell imaging showing CAVIN1-GFP signal at the PM in untreated cell, see full movie in Video S4. (F) The number of CAVIN1-GFP puncta at the PM in untreated HeLa cell over 120 seconds.

**A**

**Caveola associated proteins**

| Gene name | Fold change |
|---|---|
| CAVIN1; PTRF | -7.79600 |
| CAV1 | -2.30773 |
| EHD1 | -0.18238 |
| EHD4 | -0.02146 |
| EHD2 | NaN |
| CAVIN3; PRKCDBP | NaN |
| LGALS3 | 0.71335 |
| PACSIN3 | 0.79090 |
| DNM2 | 1.30874 |
| PACSIN2 | 1.62910 |

**B**

**EIF proteins**

| Gene name | Fold change |
|---|---|
| EIF6 | -0.25637 |
| EIF4A3 | -0.14103 |
| EIF4G3 | NaN |
| EIF4E2 | NaN |
| EIF1B | NaN |
| EIF2B5 | NaN |
| EIF2B1 | NaN |
| EIF1AD | NaN |
| EIF2B3 | NaN |
| EIF4ENIF1 | NaN |
| EIF2AK4 | NaN |
| EIF2AK2 | 0.80514 |
| EIF2S2 | 0.89536 |
| EIF4E | 1.10443 |
| EIF1 | 1.27569 |
| EIF3M | 1.32607 |
| EIF4H | 1.35615 |
| EIF1AX | 1.37635 |
| EIF4A2 | 1.42708 |
| EIF2S3 | 1.43218 |
| EIF5B | 1.45292 |
| EIF2S1 | 1.46316 |
| EIF3B | 1.49574 |
| EIF2B2 | 1.51931 |
| EIF5 | 1.52640 |
| EIF4G1 | 1.53735 |
| EIF4G2 | 1.54723 |
| EIF3F | 1.62248 |
| EIF4A1 | 1.71179 |
| EIF3E | 1.74034 |
| EIF3D | 1.80914 |
| EIF3C | 1.81512 |
| EIF3L | 1.82571 |
| EIF2A | 1.83396 |
| EIF2B4 | 1.94953 |
| EIF3I | 1.97471 |
| EIF3K | 2.01340 |
| EIF3G | 2.03053 |
| EIF3H | 2.06286 |
| EIF2D | 2.06504 |
| EIF3A | 2.06955 |
| EIF5A | 2.31074 |
| EIF3J | 3.33808 |
| EIF4B | 3.89070 |

**C**

**RPS proteins**

| Gene name | Fold change |
|---|---|
| RPS28 | -2.30421 |
| RPS27 | -1.51060 |
| RPS29 | -1.01268 |
| RPS19BP1 | -0.49721 |
| RPS15 | -0.42590 |
| RPS5 | -0.02000 |
| RPS6KA4 | NaN |
| RPS6KA3 | NaN |
| RPS23 | NaN |
| RPS6KA1 | NaN |
| RPS27L | NaN |
| RPS7 | 0.21350 |
| RPS15A | 0.56999 |
| RPS13 | 0.57028 |
| RPS9 | 0.69977 |
| RPS26 | 0.75215 |
| RPS8 | 0.75759 |
| RPS3A | 0.75787 |
| RPS11 | 0.80710 |
| RPS24 | 0.84268 |
| RPS17 | 0.92381 |
| RPS10 | 0.93298 |
| RPS14 | 0.93831 |
| RPS6 | 0.96423 |
| RPS27A | 0.98486 |
| RPS4X | 0.98834 |
| RPS16 | 1.13091 |
| RPS2 | 1.18173 |
| RPS19 | 1.22810 |
| RPS25 | 1.42511 |
| RPS3 | 1.50987 |
| RPS18 | 1.51218 |
| RPS20 | 1.51355 |
| RPS12 | 1.80088 |
| RPSA | 1.99897 |
| RPS21 | 2.21402 |

**D**

**RPL proteins**

| Gene name | Fold change |
|---|---|
| RPL35A | -2.15668 |
| RPL31 | -1.82624 |
| RPL7L1 | -0.19274 |
| RPL6 | -0.14041 |
| RPL12 | -0.04234 |
| RPL28 | NaN |
| RPL29 | NaN |
| RPL26 | NaN |
| RPL39P5 | NaN |
| RPL36AL;RP | NaN |
| RPL22L1 | NaN |
| RPL7 | 0.01322 |
| RPL7A | 0.04745 |
| RPL35 | 0.07812 |
| RPL34 | 0.13344 |
| RPL18A | 0.13689 |
| RPL27A | 0.14827 |
| RPL18 | 0.15799 |
| RPL15 | 0.25596 |
| RPL3 | 0.26996 |
| RPL13A | 0.34264 |
| RPL26L1 | 0.34612 |
| RPL17 | 0.40207 |
| RPLP0 | 0.42359 |
| RPL13 | 0.42733 |
| RPL22 | 0.44704 |
| RPL30 | 0.45132 |
| RPLP2 | 0.46483 |
| RPL27 | 0.47930 |
| RPL10A | 0.48813 |
| RPL32 | 0.49748 |
| RPL14 | 0.56716 |
| RPL36 | 0.58011 |
| RPL4 | 0.61497 |
| RPL23A | 0.62162 |
| RPL19 | 0.69175 |
| RPL8 | 0.70836 |
| RPL23 | 0.71591 |
| RPL21 | 0.74240 |
| RPLP1 | 0.77767 |
| RPL24 | 0.82356 |
| RPL37A | 0.83806 |
| RPL9 | 0.89411 |
| RPL5 | 0.93282 |
| RPL10 | 1.00989 |
| RPL11 | 1.02409 |
| RPL38 | 1.37588 |

**E**

**tRNA ligases**

| Gene name | Fold change |
|---|---|
| HARS2 | -1.37556 |
| MARS2 | -1.34851 |
| PARS2 | -1.16353 |
| RARS2 | -1.14075 |
| AARS2 | -1.11095 |
| EARS2 | -1.03670 |
| TARS2 | -0.96683 |
| VARS2 | -0.75183 |
| HARS | -0.70910 |
| CARS2 | -0.51657 |
| NARS2 | -0.37715 |
| SARS2 | -0.11476 |
| LARS2 | -0.05611 |
| IARS2 | -0.02943 |
| WARS2 | 0.09191 |
| YARS2 | 0.11797 |
| GARS | 0.13489 |
| DARS2 | 0.81510 |
| FARS2 | 0.85224 |
| FARSB | 1.22242 |
| KARS | 1.32401 |
| YARS | 1.63208 |
| FARSA | 1.65773 |
| AARS | 1.75609 |
| CARS | 1.77796 |
| LARS | 1.84042 |
| RARS | 1.90521 |
| IARS | 1.98489 |
| DARS | 2.00845 |
| MARS | 2.04212 |
| NARS | 2.12335 |
| QARS | 2.13369 |
| SARS | 2.14680 |
| EPRS | 2.53212 |
| WARS | 2.75939 |
| TARS | 2.96964 |
| VARS | 4.87209 |

Figure S6

**Figure S6.** Lists of dysregulated proteins in CAVIN1-null cell line that belong to caveolar associated proteins (A), EIF proteins (B), RPS proteins (C), RPL proteins (D) and tRNA ligases (E).

**Table S1.** Comparative proteomics revealed changes in proteins levels in CAVIN1-null HeLa cells. Fold changes in proteins levels are indicated in column O ("Difference $Log_2$ LFQ int. Cavin1KO/HeLaWT).

**Table S2.** Pathway analysis and upstream regulator analysis on dysregulated proteins in CAVIN1-null HeLa cells. QIAGEN IPA-identified upregulated pathways, downregulated pathways, upstream regulators and downregulated canonical pathways in CAVIN1-null cells are presented as the first, second, third and fourth worksheet respectively in the excel file.

**Supplementary Text**

Pathway analysis revealed a major role for CAVIN1 in protein translation (Table S2), specifically proteins identified as EIFs (eukaryotic initiation factors) (31/44) (Figure S6B) and tRNA ligases (18/36) (Figure S6E) and were significantly upregulated. In addition, proteins of the ribosomal small subunit (RPS) (26/36) (Figure S6C) and ribosomal large L subunit (RPL) (37/46) (Figure S6D) were mostly increased in CAVIN1-null cells. Other pathways that were significantly upregulated in *CAVIN1* KO cells include a number of proteins involved in DNA replication such as mini-chromosome maintenance complex-binding protein (MCMBP) (3.2-fold) and the nucleotide excision repair protein homolog MMS19 (1.9-fold), and several proteins broadly involved in stress responsive pathways such TKT (3.3-fold) and PRDX6 (2.9-fold).

CAVIN1 was originally described as a nuclear protein regulating ribosomal RNA transcription[80]. Recent studies in adipocytes demonstrated that insulin can cause translocation of CAVIN1 to the nucleus that requires CAVIN1 phosphorylation[19] in a pathway that promotes ribosomal RNA transcription. This pathway was demonstrated to be necessary for increased ribosomal biogenesis in response to growth factor stimulation. Loss of CAVIN1 caused an increase in ribosomal protein production that resulted in p53-mediated ribosomal stress. Other studies have also identified a requirement for CAVIN1 in oxidant-induced sequestration of MDM2, a negative regulator of p53, into caveolar membranes away from p53 to allow activation of the p53/p21 pathway[81]. In support of these findings, our whole proteomic analysis of CAVIN1-null HeLa cells also suggests intimate involvement of CAVIN1 in ribosomal RNA transcription through the regulation of RPS and RPL ribosomal proteins, EIFs and tRNA ligases, and is indicative of nucleolar/ribosomal stress in these cells under steady state conditions.

Ribosomal biogenesis also causes substantial demands on the nuclear import and export pathways. As a result, perturbation of nuclear import/export can also elicit ribosomal biogenesis stress. Interestingly, CAVIN1-null cells specifically exhibited an increase of a number of import proteins including importin (IPO)-4 (3.5-fold) and IPO7 (1.3-fold). Knockdown of IPO7 has been shown to cause an imbalance in the nuclear import of ribosomal proteins which is sufficient to trigger RPL5- and RPL11-dependent p53 activation, confirming that it occurs as a consequence of ribosome biogenesis stress.

Compared to the number of upregulated proteins, we noted that fewer proteins were downregulated in CAVIN1-null cells (Figure 1A; Table S1), contributing to the dysregulation of canonical pathways including caveola mediated endocytosis signalling, cholesterol biosynthesis I, cholesterol biosynthesis

II via 24, 25 dihydrolyolanosterol, cholesterol biosynthesis III via desmosterol and antigen presentation pathway (Table S2). These pathways are not surprising given the intimate relationship between caveolae and cholesterol (reviewed by Parton et al.[1]). Proteins involved in cholesterol synthesis include methylsterol monooxygenase 1 (MSMO1) (4.0-fold), squalene monooxygenase (SQLE) (2.8-fold), squalene synthase (FDFT1) (2.3-fold) and delta (24)-sterol reductase (DHCR24) (1.7-fold). This suggests that CAVIN1-null cells that lack caveolae, also have a reduced requirement for cholesterol and consequently, cholesterol biosynthesis is also decreased. Proteins linked to caveola mediated endocytosis include a number of integrins, integrin alpha-2 (ITGA2) (5.8-fold), ITGA5 (2.2-fold), ITGA3 (2.1-fold) and ITGA6 (1.5-fold). Downregulation of CAV1 has been demonstrated to lead to a reduction in integrin endocytosis that further reduced fibronectin matrix turnover[82,83]. In addition, CAVIN1 has also been implicated as a potent inhibitor of the clathrin-independent carriers/GPI-AP enriched endosomal compartment (CLIC/GEEC) endocytic pathway, in a process that is independent of caveolae formation where it was shown that noncaveolar CAVIN1 can act on cell division control 42 homolog (CDC42), a key regulator of this pathway[84]. CDC42 is also downregulated in CAVIN1-null cells (0.85-fold). Collectively, these findings suggest that CAVIN1 is intimately involved in caveola mediated endocytosis and cholesterol biosynthesis.
